# The PI3K-AKT-mTOR axis persists as a therapeutic dependency in KRAS^G12D^-driven non-small cell lung cancer

**DOI:** 10.1101/2023.09.20.558592

**Authors:** W. J. McDaid, L. Wilson, H. Adderley, M. J. Baker, J. Searle, L. Ginn, T. Budden, M. Aldea, A. Marinello, J. Aredo, A. Viros, B. Besse, H. A. Wakelee, F. Blackhall, C. R. Lindsay, A. Malliri

## Abstract

**Introduction:** KRAS^G12C^ and KRAS^G12D^ inhibitors represent a major translational breakthrough for non-small cell lung cancer (NSCLC) and cancer in general by directly targeting its most mutated oncoprotein. However, resistance to these small molecules has highlighted the need for rational combination partners necessitating a critical understanding of signaling downstream of KRAS mutant isoforms.

**Methods:** We contrasted tumor development between *Kras^G12C^*and *Kras^G12D^* genetically engineered mouse models (GEMMs). To corroborate findings and determine mutant subtype-specific dependencies, isogenic models of *Kras^G12C^* and *Kras^G12D^* initiation and adaptation were profiled by RNA sequencing. We also employed cell line models of established KRAS mutant NSCLC and determined therapeutic vulnerabilities through pharmacological inhibition. We analysed differences in survival outcomes for patients affected by advanced *KRAS^G12C^* or *KRAS^G12D^*-mutant NSCLC.

**Results:** KRAS^G12D^ exhibited higher potency *in vivo*, manifesting as more rapid lung tumor formation and reduced survival of KRAS^G12D^ GEMMs compared to KRAS^G12C^. This increased potency, recapitulated in an isogenic initiation model, was associated with enhanced PI3K-AKT-mTOR signaling. However, KRAS^G12C^ oncogenicity and downstream pathway activation were comparable with KRAS^G12D^ at later stages of tumorigenesis *in vitro* and *in vivo*, consistent with similar clinical outcomes in patients. Despite this, established KRAS^G12D^ NSCLC models depended more on the PI3K-AKT-mTOR pathway, while KRAS^G12C^ models on the MAPK pathway. Specifically, KRAS^G12D^ inhibition was synergistically enhanced by AKT inhibition.

**Conclusions:** Our data highlight a unique combination treatment vulnerability and suggest that patient selection strategies for combination approaches using direct KRAS inhibitors should be i) contextualised to individual RAS mutants, and ii) tailored to their downstream signaling.

## Introduction

Non-small cell lung cancer (NSCLC) is the most common form of lung cancer, diagnosed in up to 85% of patients^1^. It is the leading cause of cancer-related deaths worldwide, with only ∼20% of NSCLC patients surviving longer than 5 years^2,3^. Lung adenocarcinoma (LUAD), originating in alveolar type 2 epithelial cells, is the most common histological subtype of NSCLC^4^, with KRAS the most frequently mutated oncogenic driver, found in ∼30% of cases^5^. KRAS gain-of-function mutations are critical for both the initiation and maintenance of tumors^6^, with different KRAS point mutations occurring with varying prevalence. KRAS^G12C^ is the most common mutation occurring in up to 40% of KRAS-mutant LUAD, followed by KRAS^G12V^ and KRAS^G12D^, which occur in up to 19% and 15% of KRAS-mutant LUAD, respectively^7^.

It is now accepted that NSCLC bearing different KRAS mutations are heterogenous resulting from factors such as varying levels of KRAS activation (GTP-KRAS), upregulation of distinct pro-tumorigenic functions and contextual acquisition of secondary mutations exclusive to each KRAS mutation^8–11^. Consequently, a ‘one drug fits all’ approach to targeting KRAS mutant lung cancer is challenging, and treatment should be tailored to the subtype of KRAS mutation. This is exemplified through clinical trial disappointments such as the failure of MEK inhibitors to show meaningful improvements to patients with KRAS-mutant NSCLC in the SELECT-1 study, heavily implying that a greater resolution of precision medicine is required for RAS targeting^12^. For this to be achieved, it is important to characterise the signaling pathways activated by different KRAS mutants and identify dependencies exclusive to each mutant isoform that can be exploited therapeutically.

In recent years, direct inhibitors of KRAS^G12C^ and KRAS^G12D^ have been developed with KRAS^G12C^ inhibitors, such as Sotorasib and Adagrasib, now in the clinic, with the former inducing responses in 25-40% of patients^13–17^. However, the efficacy of KRAS^G12C^ inhibitors is limited by several intrinsic resistance mechanisms^18^, also expected to impede KRAS^G12D^ inhibitors that are currently being assessed in early-phase clinical trials. In addition, it is now known that KRAS^G12D^ is associated with immune suppression and resistance to PD-L1 therapy compared to other KRAS mutant isoforms^19–21^. It is therefore critical to identify KRAS^G12D^-specific dependencies to target in combination with KRAS^G12D^ inhibition, aiming to improve patient outcomes by overriding anticipated resistance.

In this study, through a comprehensive analysis of isogenic systems with validation in physiologically relevant tumor cell models and NSCLC patient data, we investigated the biological features of KRAS^G12C^ and KRAS^G12D^ mutations and their signaling differences during NSCLC initiation and in established cell models. We identified a KRAS^G12D^-specific mechanism of tumorigenesis which is therapeutically exploitable and potentiates KRAS^G12D^ inhibition.

## Materials and Methods

### In vivo studies

All mouse studies were carried out in compliance with UK Home Office regulations with protocols approved by the Cancer Research UK Manchester Institute Animal Welfare and Ethical Review Advisory Body. Generation of the *Kras^G12C^* mouse model, tumor burden, survival and early lesion studies, and histological analyses are described in Supplementary Materials and Methods. Sequences for gRNA, repair template ultramer and genotyping primers are described in Supplementary Table S1.

### Cell culture

All cell lines were cultured at 37°C with 5% CO_2_. H358, HCC1171, H1792, H2030, H23, HOP62, A427, SKLU-1 and HCC461 cells were cultured in RPMI-1640 Medium (Gibco, #21875034) supplemented with 10% FBS (Biosera, #FB-1001T) and 1% penicillin/streptomycin (P/S) (Gibco, #15140122). MEFs and Lenti-X 293T cells were cultured in DMEM (Gibco, #41966-029) with 10% FBS and 1% P/S. MEFs were maintained in 4µg/mL blasticidin to maintain expression of KRAS transgenes. GEMM-derived tumor cell lines were cultured in DMEM/F12 (Gibco, #11330-032) supplemented with 10% FBS, 1% P/S, 2mM glutamine, 1µM hydrocortisone, 20ng/mL murine EGF (Cell Signaling, #5331) and 50 ng/mL murine IGF (Bio-techne, #791-MG-050). MLE-12 cells were cultured in DMEM/F12 supplemented with 2% FBS, 1% P/S, 2nM glutamine, 5µg/mL insulin, 10µg/mL transferrin, 30nM sodium selenite, 10nM hydrocortisone, 10mM HEPES and 10nM β-estradiol. For experiments, MLE-12 cells were cultured in the same media but with 0.5% FBS. Lenti-X 293T cells were purchased from Takarabio. H358, H23, A427, SKLU-1, HCC1171, H1792, H2030, HOP62 and MLE-12 cells were purchased from ATCC. HCC461 cells were kindly donated by John Minna (UT Southwestern, USA). KRAS MEFs were provided by Frederick National Lab for Cancer Research (NCI, USA). Cell lines were regularly authenticated and checked for mycoplasma contamination through in-house facilities. Generation of cell models, proliferation and viability assays, Western blotting, RNA sequencing and analysis, intracellular and surface staining by flow cytometry, caspase-3/-7 detection and propidium iodide staining are described in Supplementary Materials and Methods. Antibodies are documented in Supplementary Table S2.

### Clinical database analysis

575 RAS mutant patients were recruited to the multicentre transatlantic RAS precision medicine (RAS-PM) study from three tertiary cancer centres including The Christie NHS Foundation Trust, The Gustave Roussy Cancer Centre and The Stanford Cancer Institute. Key inclusion criteria were defined as: approval by the ethics committees as required by local or international standards, stage IIIb/IV NSCLC, availability of progression-free survival (PFS) and overall survival (OS) data and confirmed RAS mutant status. Key exclusion criteria included: inconclusive or no confirmed NSCLC diagnosis histologically, patients with cancers wild-type for KRAS and other mutations other than KRAS^G12C^ and KRAS^G12D^, and no PFS or OS data. First-line PFS was defined as time from treatment start, in advanced stage disease, to progression or death from any cause. OS is defined as time from first-line treatment start, in advanced stage disease, to death regardless of cause. Patients still alive at last visit are censored at date of last follow-up. Data collection protocols were approved by local governance committee.

### Statistical analysis

All error bars shown on graphs represent ± standard error of the mean (s.e.m.). The specific statistical tests used are indicated in the figure legends alongside the p values and were carried out using GraphPad Prism 10. For comparison between two conditions, statistical tests can be assumed to be a two-tailed Student’s t-test. For multiple comparisons, one-way or two-way ANOVA was used. Log-rank test was used for survival curve analyses. n.s. = not significant, *P< 0.05, **P< 0.01, ***P< 0.001 and **** P < 0.0001.

## Results

### KRAS^G12D^ is more potent than KRAS^G12C^ in driving NSCLC initiation in vivo

To examine and compare the oncogenic potency of *Kras^G12D^*and *Kras^G12C^ in vivo*, we used a well-characterized genetically-engineered mouse model (GEMM) that harbours latent *Kras^G12D^*, whose activation with adenovirus expressing Cre recombinase (AdVC) drives the formation of lung tumors closely resembling human LUAD^23^ (Figure 1A,B). We then used CRISPR/Cas9 technology to convert the aspartic-acid-encoding codon 12 to one encoding cysteine (*Kras^G12C^*) to create a *Kras^G12C^* mouse model (Figure 1B). Thus, we could study relative potency of *Kras^G12D^* and *Kras^G12C^* in a closely controlled *in vivo* setting. We further combined both *Kras* alleles with conditional *tp53* knockout (*tp53^KO^*) to accelerate tumorigenesis and better recapitulate human NSCLC (Figure 1B)^24^.

**Figure 1.**
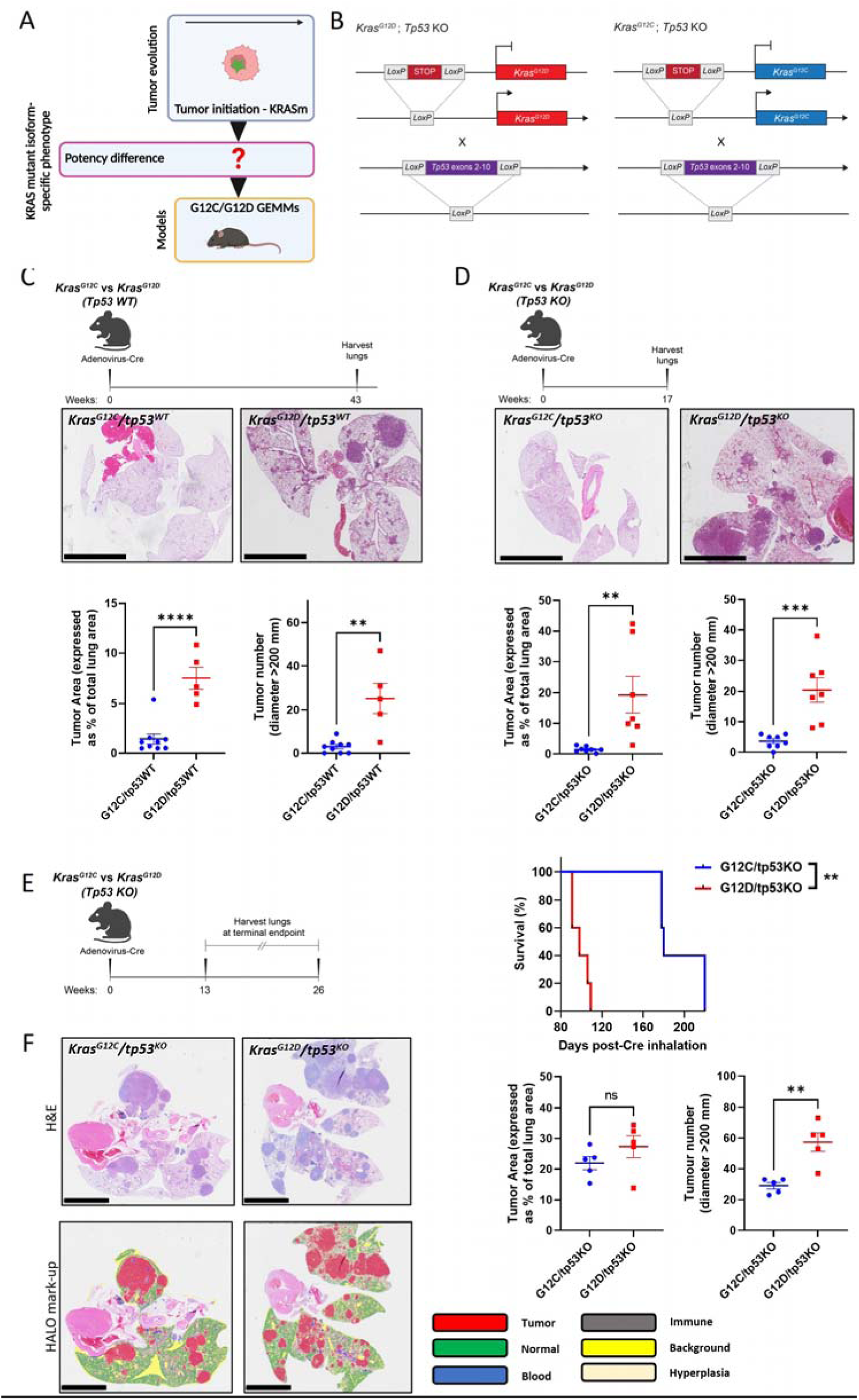
KRAS^G12D^ is more potent than KRAS^G12C^ in driving NSCLC initiation *in vivo*. (A) Schematic illustrating the use of GEMM models to study the impact of KRAS mutant isoforms on NSCLC initiation. Green star = KRAS mutation. Created with BioRender.com. (B) Schematic illustrating KRAS mutant oncogenes silenced by the insertion of a STOP codon flanked by LoxP sites. AdVC administration by inhalation leads to LoxP site recombination, removing the STOP codon allowing KRAS mutant isoform expression in GEMM mice lungs. Conditional KRAS mutant mice were crossed with mice in which *tp53* is also flanked by LoxP sites. AdVC induces LoxP recombination and loss of p53 protein expression. (C) (Above) Timeline of experiment. (Below) Representative H&E sections and HALO quantification of lung tumor area and number per mouse comparing *Kras^G12C^/tp53^WT^* and *Kras^G12D^/tp53^WT^* mice 11 months after AdVC exposure (n=10 *Kras^G12C^/tp53^WT^* mice and 5 *Kras^G12D^/tp53^WT^*mice); scale bar = 5mm. (D) (Above) Timeline of experiment. (Below) Representative H&E sections and HALO quantification of lung tumor area and number comparing *Kras^G12C^/tp53^KO^* and *Kras^G12D^/tp53^KO^*mice 4 months after AdVC exposure (n= 8 mice per genotype); scale bar = 5mm. (E) (Left) Timeline of experiment. (Right) Survival analysis for *Kras^G12C^/tp53^KO^* and *Kras^G12D^/tp53^KO^*mice after intranasal delivery of AdVC (n=5 mice per genotype, Log-Rank (Mantel-Cox) test). (F) (Top left) Representative H&E images, (Bottom left) HALO mark-up and (Right) HALO quantification of tumor area and number comparing *Kras^G12C^/tp53^KO^* and *Kras^G12D^/tp53^KO^*mice from survival study (n=5 mice per genotype); scale bar = 5mm. C,D and F depict mean ±s.e.m and statistical analysis carried out using unpaired Student’s t-test. ****P<0.0001, **P<0.01, ns>0.05.

*Kras^G12C^* mouse models have so far been under-reported in NSCLC research, highlighting the value this tool offers to the investigation of lung cancers driven by this oncogene. We first confirmed that the *Kras^G12C^* mouse model was functional: after AdVC inhalation, lung tumors were formed which recapitulated tumors resembling lung adenocarcinoma, similarly to the *Kras^G12D^* mouse model (Supplementary Figure S1A). However, when *Kras^G12C^* were compared with *Kras^G12D^* mice matched for time after AdVC inhalation, there was a striking difference in tumorigenic properties between *Kras^G12D^* and *Kras^G12C^* models on both *tp53* wild-type (*tp53^WT^*) and *tp53^KO^* backgrounds: *Kras^G12D^*-expressing mice had a dramatically increased tumor burden compared to *Kras^G12C^* (Figure 1C,D). *Kras^G12D^* GEMMs also had increased hyperplasia compared to *Kras^G12C^* GEMMs (Supplementary Figure S1B,S1C). This difference in tumor burden was reflected by shorter median overall survival of *Kras^G12D^/tp53^KO^* GEMMs (98 days) compared to *Kras^G12C^/tp53^KO^* GEMMs (180 days) (Figure 1E). These results were reproduced using an alternative method of virus inhalation with similar tumor latency observed (median survival was 211 days for *Kras^G12C^/tp53^KO^* mice vs 102 days for *Kras^G12D^/tp53^KO^* mice (Supplementary Figure S1D). However, despite a slight non-significant increase in lung tumor area for *Kras^G12D^/tp53KO* and comparable hyperplasia areas between *Kras^G12C^/tp53^KO^*and *Kras^G12D^/tp53^KO^* mice sacrificed due to disease symptoms (Figure 1F and Supplementary Figure S1E), *Kras^G12D^* mice had approximately two-fold higher tumor number, constituted mainly by an increased number of smaller tumors (Figure 1F and Supplementary Figure S1F). These data suggest that *Kras^G12D^* is more effective than *Kras^G12C^*in initiating NSCLC tumors.

### KRAS^G12D^ co-opts the PI3K-AKT-mTOR pathway to promote tumor initiation in NSCLC

We next asked whether the increased oncopotency of *Kras^G12D^*compared to *Kras^G12C^* may be due to signaling differences immediately downstream upon tumor initiation. To investigate this, we generated an isogenic KRAS mutant initiation model using an immortalised murine lung alveolar type 2 cell line, MLE-12, that is non-tumorigenic when inoculated in mice^25^, and modified it to ectopically express either flag-tagged wildtype KRAS (KRAS^WT^), KRAS^G12C^ or KRAS^G12D^ under the control of a doxycycline-regulated promoter (Figure 2A,B). We confirmed the correct expression of each isoform through exposure of the isogenic panel to doxycycline in the presence or absence of the KRAS^G12C^ inhibitor (G12Ci) Sotorasib. Exposure to G12Ci only affected *Kras^G12C^* MLE-12 cells by binding to KRAS and switching off MAPK signaling as evidenced by reduced phosphorylated ERK. Additionally, we were able to detect KRAS^G12D^ protein expression using a KRAS^G12D^-specific antibody only in *Kras^G12D^* MLE-12 cells (Supplementary Figure S2A). MLE-12 cells were next cultured in ultra-low attachment (ULA) plates to induce growth as 3D spheroids, mimicking more closely the physical characteristics of cancer cells in a tumor^26^. Upon doxycycline treatment, an increase in metabolic activity and spheroid size was observed in *Kras^G12D^*-compared to *Kras^G12C^*-initiated cells, indicative of increased proliferation and consistent with our *in vivo* data (Figure 2C and Supplementary Figure S2B).

**Figure 2.**
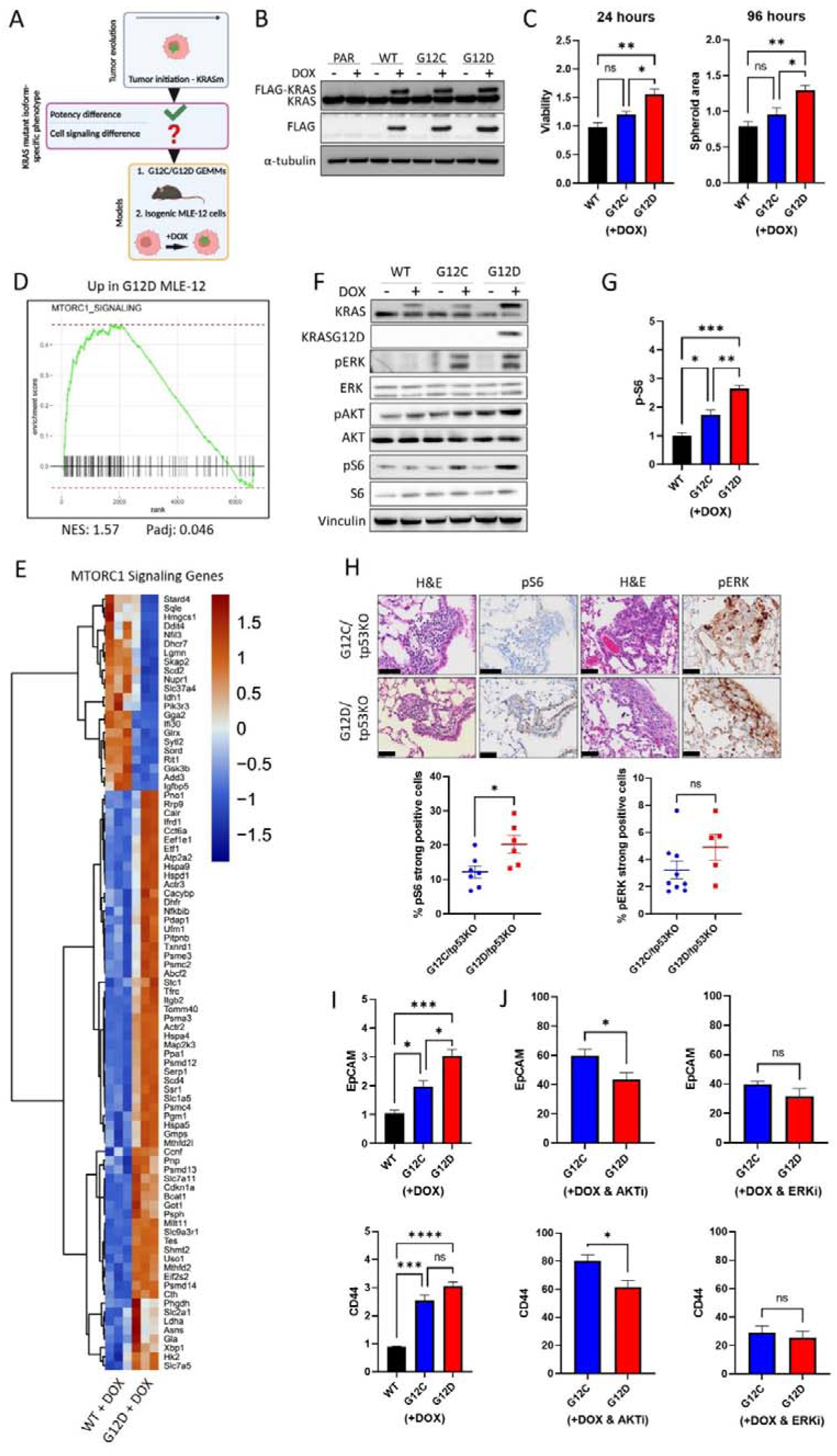
KRAS^G12D^ co-opts the PI3K-AKT-mTOR pathway to promote tumor initiation in NSCLC. (A) Schematic illustrating the use of GEMM models and isogenic MLE-12 cells to study the impact of KRAS mutant isoforms on NSCLC initiation and signaling differences between isoforms. Green star = KRAS mutation. Created with BioRender.com. (B) Western blot analysis of KRAS and FLAG-tagged KRAS upon 24 hour exposure of isogenic MLE-12 cells to 100ng/mL doxycycline. PAR = parental. (C) (Left) MLE-12 spheroid viability upon 24 hour 100ng/mL doxycycline exposure measured by CellTiter-Glo 3D and (Right) MLE-12 spheroid area upon 96 hour 100ng/mL doxycycline exposure measured by ImageJ. Data normalised to untreated (no doxycycline) control (n=3 at 24 hours and n=4 at 96-hours). (D) GSEA showing that mTORC1 signaling genes are positively correlated with KRAS^G12D^ expression compared to KRAS^WT^. (E) Heatmap showing DEGs belonging to MTORC1 signaling gene-set when comparing *KRAS^G12D^* to *KRAS^WT^* MLE-12 cells 24 hours after 100ng/mL doxycycline exposure (n=3). (F) Western blot analysis of ERK, AKT and S6 phosphorylation 24 hours after exposure of isogenic MLE-12 cells to 100ng/mL doxycycline. Data representative of 3 independent experiments. (G) Flow cytometric quantification of S6 phosphorylation levels upon 24 hour exposure of isogenic MLE-12 cells to 100 ng/mL doxycycline. Data normalised to untreated (no doxycycline) control (n=3). (H) (Above) Representative immunohistochemical staining and (Below) quantification of ERK and S6 phosphorylation in early lung lesions of *Kras^G12C^/tp53^KO^*and *Kras^G12D^/tp53^KO^* mice (n=6-9 mice per genotype); scale bar = 50µm. (I) Flow cytometric quantification of CD44 and EpCAM surface expression in isogenic MLE-12 cells upon 24 hour exposure to 100ng/mL doxycycline. Data normalised to untreated (no doxycycline) control (n=3). (J) Flow cytometric quantification of CD44 and EpCAM surface expression in isogenic MLE-12 cells upon 24 hour exposure to 100ng/mL doxycycline in the presence of (Left) 1µM AKTi or (Right) 1µM ERKi. Data normalised to DMSO control (n=4). C,G,I and J depict mean ±s.e.m and statistical analysis carried out using one-way ANOVA test. H depicts mean ±s.e.m and statistical analysis carried out using unpaired Student’s t-test. ****P<0.0001, ***P<0.001, **P<0.01, *P<0.05, ns>0.05. DOX=doxycycline

We next asked what could be driving the increased proliferation of *Kras^G12D^*-initiated cells. Gene expression profiling of the isogenic MLE-12 panel comparing either *Kras^G12D^* or *Kras^G12C^* to *Kras^WT^* 24 hours after doxycycline exposure revealed that *Kras^G12D^* had a stronger impact on the transcriptome with 6664 differentially expressed genes (DEGs) compared to *Kras^WT^*, whereas 4566 genes were altered between *Kras^G12C^* and *Kras^WT^*. From these gene changes, we identified mTORC1 signaling as a significantly enriched gene-set when comparing *Kras^G12D^* to *Kras^WT^*(Figure 2D,E), not seen when comparing *Kras^G12C^* to *Kras^WT^*, suggesting higher activation of this pathway in cells initiated with *Kras^G12D^*. Alternatively, by comparing each isogenic cell line after addition of doxycycline to its untreated counterpart, there was a strikingly higher number of differentially expressed genes belonging to the MTORC1 signaling gene-set in cells with *Kras^G12D^* expression compared to cells with *Kras^WT^* or *Kras^G12C^*(Supplementary Figure S2C). Taken together, *Kras^G12D^* expression leads to a more extensive transcriptional reprogramming, upregulating several MTORC1-associated genes compared to *Kras^G12C^* or *Kras^WT^* MLE-12 cells.

As mTORC1 is part of the PI3K-AKT-mTOR axis, a key effector pathway of RAS signaling^27^, we postulated that KRAS^G12D^ co-opts the PI3K-AKT-mTOR pathway to a higher extent than KRAS^G12C^ and aimed to explore this pathway further as a potential mechanism underpinning the greater potency of KRAS^G12D^. Firstly, we analysed if the PI3K-AKT-mTOR pathway was hyperactivated in G12D-initiated cells. Indeed, higher phosphorylation of the mTORC1 activator, AKT, and the mTORC1 substrate ribosomal S6, were evident in *Kras^G12D^* MLE-12 cells 24 hours after exposure to doxycycline, consistent with our gene expression data (Figure 2F,G). Next, using ERK phosphorylation as a measure of MAPK signaling, we observed no difference in MAPK signaling between the two mutant isoforms (Figure 2F). By further analysing our gene expression data, we saw that DEGs related to MAPK signaling were similar between mutant isoforms (Supplementary Figure S2D). Treatment of parental MLE-12 cells with doxycycline confirmed that doxycycline does not affect either of these pathways (Supplementary Figure S2E), nor expression of exogenous KRAS^WT^ to levels matching mutant KRAS (Figure 2F,G). Higher S6 phosphorylation was also noted in early *Kras^G12D^*lung lesions *in vivo* compared to *Kras^G12C^* lesions, whilst there was no significant difference in the level of ERK phosphorylation between the two mutant isoforms (Figure 2H). Finally, we selected CD44 and EpCAM as two markers of NSCLC initiation^28–30^ and showed that their expression is increased upon induction of the mutant isoforms only (Figure 2I). Inhibition of AKT reduced expression of these markers to a greater extent in *Kras^G12D^* MLE-12 cells (Figure 2J), whereas inhibition of ERK reduced their expression to comparable levels in both cell lines. This suggests that the two mutant isoforms require input from the MAPK pathway for tumor initiation to the same extent, whereas the PI3K-AKT-mTOR pathway is required more during *Kras^G12D^*-driven initiation. Overall, these data confirmed our gene expression data and highlighted a possible role for the PI3K-AKT-mTOR pathway in tumor initiation by *Kras^G12D^*.

### Long-term KRAS^G12D^-exposed cells display specific PI3K-AKT-mTOR pathway dependency

Having established the allele-specific role of *Kras^G12D^* in facilitating tumor initiation and uncovering its increased activation of the PI3K-AKT-mTOR pathway compared to *Kras^G12C^*, we next sought to explore differences between KRAS^G12C^ and KRAS^G12D^ in advanced disease to determine if growth characteristics and signaling differences are maintained during tumor evolution (Figure 3A). As KRAS-mutant NSCLC tumors are extremely heterogenous, largely due to the diversity of their genetic alterations^9^, comparing KRAS-mutant phenotypes using patient samples or NSCLC cell lines is challenging. Therefore, we began by using a panel of isogenic MEFs, initially engineered to become ‘Ras-less’^31^ and further genetically modified to express either KRAS^WT^, KRAS^G12C^ or KRAS^G12D^ ^32^. Therefore, we could compare KRAS mutant isoforms directly without the confounding effects of co-mutations seen in lung tumors. As these cells have been cultured long-term in the presence of KRAS mutant alleles, we considered that KRAS mutant isoform-dependent adaptations have developed that may reflect features of established tumors.

**Figure 3.**
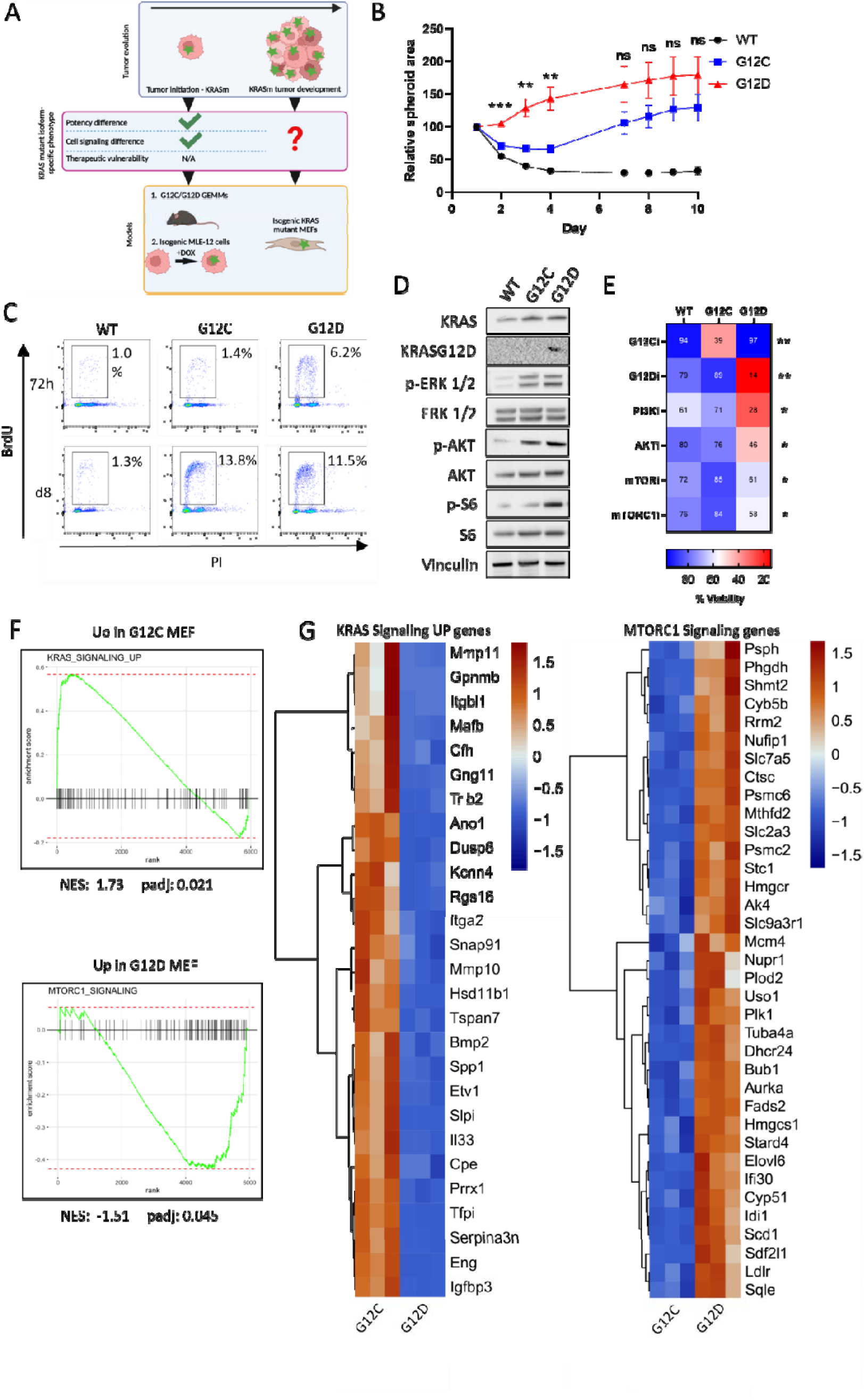
Long term KRAS^G12D^-exposed cells display specific PI3K-AKT-mTOR pathway dependency. (A) Schematic illustrating the use of isogenic KRAS MEFs to determine growth rates, signaling differences and therapeutic vulnerabilities conferred by long-term expression of KRAS mutant isoforms. Green star = KRAS mutation. Created with BioRender.com. (B) Spheroid area of isogenic KRAS MEFs quantified using ImageJ. Spheroid areas over time were normalised to spheroid area at day 1 (n=3). Statistical significance analysed using one-way ANOVA at each time point but only significance between *Kras^G12C^*and *Kras^G12D^* MEFs presented. (C) BrdU/PI staining of isogenic KRAS MEFs 72 hours and 8 days (d8) after seeding in 3D. Proliferating cells are expressed as % BrdU-positive. Data representative of three independent experiments. (D) Western blot analysis of ERK, AKT and S6 phosphorylation levels in isogenic KRAS MEFs 24 hours after seeding in 3D. Data representative of three independent experiments. (E) Viability of isogenic KRAS MEFs in response to 100nM G12Ci, 100nM G12Di, 1µM PI3Ki, 10µM AKTi and 1µM mTORi and mTORC1i in 3D. Viability was measured after 72 hours of drug exposure by CellTiter-Glo 3D. Viability expressed as % of DMSO control (n=4). Mean depicted and statistical analysis carried out using one-way ANOVA per drug treatment with significance between *Kras^G12C^* MEFs and *Kras^G12D^* MEFs presented. **P<0.01, *P<0.05 (F) GSEA showing that mTORC1 signaling genes are positively correlated with KRAS^G12D^ expression and KRAS Signaling Up genes are positively correlated with KRAS^G12C^ expression. (G) Heatmaps showing DEGs belonging to MTORC1 Signaling and KRAS Signaling UP gene-sets comparing *Kras^G12C^* to *Kras^G12D^*MEFs (n=3 per genotype).

First, we observed that *Kras^WT^* or *Kras^G12C^* MEFs were unable to proliferate when cultured in agarose, depriving cells of their anchorage. However, *Kras^G12D^* MEFs proliferated and formed colonies (Supplementary Figure S3A). To determine growth over time, we cultured the MEFs as spheroids in ULA plates and observed that *Kras^G12D^* MEFs were again able to proliferate. In contrast, *Kras^WT^* and *Kras^G12C^* MEFS were initially unable to proliferate, but over time *Kras^G12C^* MEFs began proliferating at the same rate as *Kras^G12D^*MEFs (Figure 3B,C and Supplementary Figure S3B). Similar to MLE-12 cells, ERK phosphorylation levels were comparable between *Kras^G12C^* and *Kras^G12D^* MEFs after 24 hours of culturing in 3D, while AKT and S6 phosphorylation were higher in *Kras^G12D^* MEFs (Figure 3D and Supplementary Figure S3C), suggesting that *Kras^G12D^*-specific PI3K-AKT-mTOR hyperactivation persists beyond initiation. To test if the PI3K-AKT-mTOR pathway supported anchorage-independent growth of *Kras^G12D^* MEFs, we inhibited different nodes in the pathway and observed that *Kras^G12D^*MEFs were more sensitive than *Kras^WT^* or *Kras^G12C^* MEFs (Figure 3E).

We next carried out gene expression profiling of *Kras^G12C^*and *Kras^G12D^* MEFs on day 8, aiming to determine differential signaling when both *Kras^G12C^* and *Kras^G12D^* MEFs were proliferating at the same rate. We observed that *Kras^G12C^* MEFs had increased gene expression associated with KRAS signaling (KRAS Signaling UP) whilst, similarly to *Kras^G12D^* MLE-12 cells, *Kras^G12D^* MEFs had increased expression of MTORC1 signaling genes (Figure 3F,G). Collectively, these data suggest that, in cells exposed long-term to KRAS mutant isoforms, proliferation differences may become less apparent over time, however signaling differences persist as indicated by the gene expression profiles. In order to increase oncogenicity and proliferate, KRAS^G12C^ hyperactivates KRAS signaling which may be relevant to KRAS^G12C^-driven tumor evolution. In contrast, KRAS^G12D^ relies more on PI3K-AKT-mTOR signaling, even beyond initiation, which may be therapeutically exploitable.

### KRAS^G12C^ and KRAS^G12D^ NSCLC cell lines exhibit RAS effector-specific dependencies

Our above findings from the mutant KRAS MEFs led us to hypothesise that, once KRAS^G12C^- and KRAS^G12D^-driven tumors are established, the difference in potency between the two variants becomes less evident. Given that precision medicine and KRAS inhibitors are usually administered in the context of stage IV NSCLC, we asked whether KRAS mutant-specific dependency on the pathways described above persists in established NSCLC tumors and cell lines, conferring isoform-specific vulnerabilities (Figure 4A). First, we carried out immunohistochemical (IHC) staining of tumors from the GEMM survival study (Figure 1E) for markers of proliferation (Ki67), cell cycle progression (Cyclin D1) and ERK and S6 activation. Interestingly, we did not observe significant differences in the levels of staining for these proteins between *Kras^G12C^/tp53^KO^* and *Kras^G12D^/tp53^KO^*lung tumors (Figure 4B,C) implying that there is no longer a potency (proliferation) or signaling difference between the two mutant isoforms, possibly due to acquired mutations affecting activation of these pathways^33^. In agreement, there was no significant difference in proliferation (Supplementary Figure S4) or ERK, AKT and S6 activation (Figure 4D) between murine tumor cell lines (mTCL) derived from *Kras^G12C^/tp53^KO^*and *Kras^G12D^/tp53^KO^* GEMMs, further implying that the genotype-specific difference in potency and signaling was lost in established tumors. However, *Kras^G12C^* mTCL was more sensitive to ERK and MEK inhibition, whilst *Kras^G12D^* mTCL was more sensitive to PI3K, AKT and mTOR inhibition (Figure 4E), despite both cell lines showing similar levels of MAPK and PI3K-AKT-mTOR pathway activation, implying that the vulnerabilities persist independently of phosphorylation levels.

**Figure 4.**
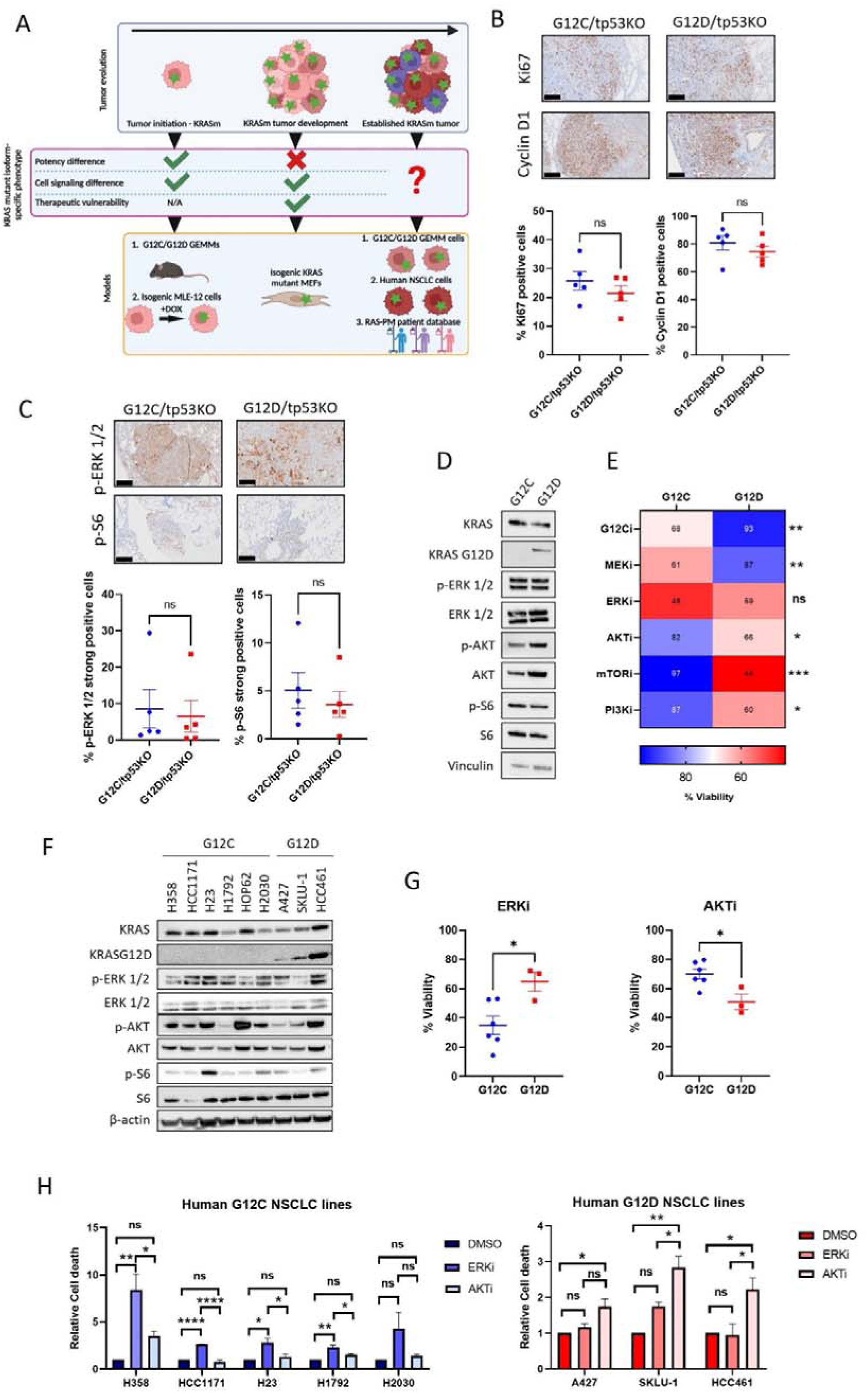
KRAS^G12C^ and KRAS^G12D^ NSCLC cells exhibit RAS effector-specific dependencies. (A) Schematic illustrating the use of GEMM-derived NSCLC cell lines, human NSCLC cell lines and patient data to determine growth rates, signaling differences and therapeutic vulnerabilities imposed by KRAS mutant isoforms in advanced disease. Green star = KRAS mutation. Created with BioRender.com. (B) (Above) Representative immunohistochemical staining and (Below) quantification of Ki67 and Cyclin D1 expression in established lung tumors of *Kras^G12C^/tp53^KO^* and *Kras^G12D^/tp53^KO^* mice (n=5 per genotype). Scale bar = 200µm. (C) (Above) Representative immunohistochemical staining and (Below) quantification of ERK and S6 phosphorylation in established lung tumors of *Kras^G12C^/tp53^KO^*and *Kras^G12D^/tp53^KO^* mice (n=5 per genotype). Scale bar = 200µm. (D) Western blot analysis of ERK, AKT and S6 phosphorylation in *Kras^G12C^* and *Kras^G12D^* mTCLs 48 hours after seeding in 3D. Data representative of three independent experiments. (E) Viability of *Kras^G12C^* and *Kras^G12D^* mTCLs in response to 1µM G12Ci, 10µM G12Di, 10µM MEKi, 10µM ERKi, 1µM PI3Ki, 10µM AKTi and 10nM mTORi. Viability was measured after 48 hours of drug exposure by crystal violet staining. Viability expressed as % of DMSO control (n=3). (F) Western blot analysis of ERK, AKT and S6 phosphorylation of *Kras^G12C^* and *Kras^G12D^* human NSCLC cell lines 48 hours after seeding in 3D. Data representative of three independent experiments. (G) Viability of human *KRAS^G12C^* and *KRAS^G12D^* NSCLC cell lines in response to 10µM ERKi and 10µM AKTi in 3D. Viability was measured after 72 hours of drug exposure by CellTiter-Glo 3D. Viability expressed as % of DMSO control (n=3). (H) Cell death analyses of human (Left) *KRAS^G12C^* and (Right) *KRAS^G12D^* NSCLC cell lines in response to 10µM ERKi and 10µM AKTi in 3D. Cell death was measured after 48 hours of drug exposure by flow cytometric quantification of PI staining. Data normalised to DMSO control (n=3). B, C, E and G depict mean ±s.e.m and statistical analysis carried out using unpaired student’s t-test. H depicts mean ±s.e.m and statistical analysis carried out using one-way ANOVA ****P<0.0001, ***P<0.001, **P<0.01, *P<0.05, ns>0.05.

To determine if these findings were also relevant to human NSCLC, we first examined differences in KRAS mutant isoform-specific survival outcomes from an internationally recruited cohort of advanced KRAS-mutant NSCLC patients, the RAS-Precision Medicine (RAS-PM) database. Of 575 patients recruited, 240 were affected by cancers harboring *KRAS^G12C^*, compared to 92 patients with cancers harboring *KRAS^G12D^* (Supplementary Figure S5A). There were no significant differences in baseline characteristics between *KRAS^G12C^* and *KRAS^G12D^* mutant patients (Supplementary Table S3). There was also no significant difference in either 1^st^ line progression-free survival (PFS) or overall survival (OS) (Supplementary Figure S5B,5C). To test whether mutant-subtype specific differences were more apparent at earlier stages of tumorigenesis in a clinical cohort, we next extracted data from cBioPortal to examine relative differences between KRAS^G12C^ and KRAS^G12D^ NSCLC (Supplementary Figure S5D)^34,35^. We analyzed data from three cohorts: NSCLC TRACERx study (2017)^36^, TCGA Firehose Legacy and Pan-Lung Cancer study (2016)^37^. In contrast to RAS-PM, this cohort was considered early stage and operable, with the intention of examining differences in RAS subtypes at point of diagnosis rather than deterioration. In line with our preclinical observations, the proportion of KRAS^G12D^ T3 and T4 stage tumors was higher compared to KRAS^G12C^ NSCLC (Supplementary Figure S5E). Taken together, these clinical results parallel our *in vitro* and *in vivo* findings, highlighting that the tumorigenic strength of KRAS^G12D^ becomes less apparent at late stages of NSCLC, whereby the oncopotency of KRAS^G12C^ appears to ‘catch up’ with KRAS^G12D^.

Despite the loss of potency, we wondered if signaling and therapeutic differences persist in advanced human disease. We selected a panel of 6 KRAS^G12C^ and 3 KRAS^G12D^ human NSCLC cell lines. We first assessed proliferation rates among these lines and saw no significant difference in proliferation, underpinning the late-stage *in vivo* and patient data (Supplementary Figure S6A). Similar to established tumours in GEMMs, there was no clear differences in ERK, AKT and S6 activation between human KRAS^G12C^ and KRAS^G12D^ cell lines (Figure 4F), again likely due to factors such as genomic heterogeneity between cell lines or Epithelial-Mesenchymal Transition influencing pathway activation^38^. However, when we mined publicly available data from the Cancer Cell Line Encyclopedia (CCLE) and compared gene expression data between *KRAS^G12C^* and *KRAS^G12D^* cell lines^39^, human *KRAS^G12C^*lines were enriched for genes related to increased KRAS signaling, while human *KRAS^G12D^* lines were enriched for genes related to PI3K-AKT-mTOR and specifically mTORC1 signaling (Supplementary Figure S6B), similarly to our MEF gene expression data. We next tested the impact of MEK, ERK and AKT inhibition in these NSCLC cell lines and observed that *KRAS^G12C^*lines were more sensitive to ERK or MEK inhibition, while *KRAS^G12D^* lines were more sensitive to AKT inhibition (Figure 4G and Supplementary Figure S6C). Additionally, we observed higher cell death after ERK or AKT inhibition in *KRAS^G12C^* or *KRAS^G12D^* cells, respectively (Figure 4H). Altogether, these data imply that, in advanced disease, the difference in potency between these two KRAS mutant isoforms is no longer apparent in terms of proliferation and immediate signaling. However, *KRAS^G12C^* and *KRAS^G12D^* cells are more susceptible to MAPK and PI3K-AKT-mTOR pathway inhibition, respectively. Therefore, these mutant-subtype specific treatment vulnerabilities persist despite the loss of clear differences in oncogenic signaling and phenotype at this point of NSCLC evolution.

The clinical development of direct KRAS^G12C^ inhibitors provides our most advanced current means of inhibiting KRAS^13^. However, the clinical efficacy of KRAS^G12C^ inhibition (G12Ci) in NSCLC is hindered by intrinsic factors such as pathway re-activation and feedback/bypass pathways which often result in resistance^18,40^. It is expected that resistance will circumvent KRAS^G12D^ inhibition (G12Di) in NSCLC, with reports of resistance mechanisms and combination strategies already emerging in colorectal cancer^41,42^. Thus, it is vital to explore potential combination therapies to minimise resistance and maximise the potential of G12Di. Having identified the PI3K-AKT-mTOR axis as a KRAS^G12D^-specific vulnerability, we next examined whether its inhibition combines effectively with G12Di in NSCLC. We found that in our isogenic MEF panel the combined effect of G12Di + AKTi was synergistic (C.I.value<1) across multiple doses of G12Di and, overall, more potent compared to that of the G12Ci + AKTi combination, where an additive effect was observed across all doses (Figure 5A and Supplementary Figure S7A). This suggested that co-targeting KRAS^G12D^ and the PI3K-AKT-mTOR axis, represents a potential drug combination. Consistent with this, using our GEMM-derived cell lines of KRAS^G12C^ and KRAS^G12D^-driven NSCLC, we again saw that G12Di + AKTi acted synergistically and was more potent compared to G12Ci + AKTi, which again was additive in terms of combination effect (Supplementary Figure S7B).

**Figure 5:**
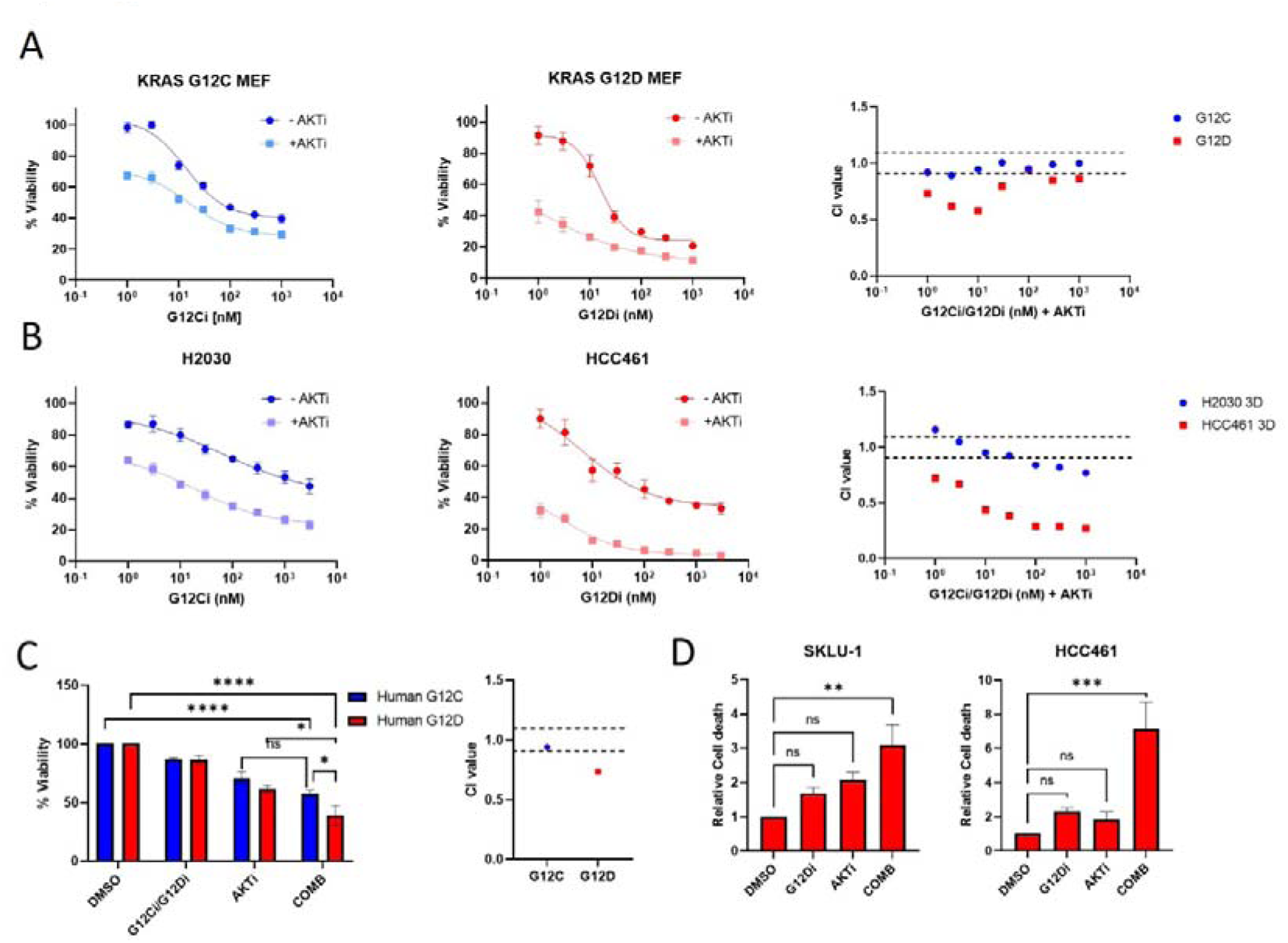
KRAS^G12D^ inhibition and PI3K-AKT-mTOR inhibition synergise in KRAS^G12D^ cells. (A) Isogenic MEFs were treated with increasing concentrations of either G12Ci or G12Di in the presence or absence of 10µM AKTi in 3D. 48 hours later, viability was measured by CellTiter-Glo 3D. Viability expressed as % of DMSO control. CI values were calculated (n=3). (B) The same conditions and analysis as 5A in H2030 (KRAS^G12C^) and HCC461 (KRAS^G12D^) NSCLC cell lines (n=3). (C) Human NSCLCs were treated with either 1nM (for H358, HOP62, H2030 and HCC1171), 5nM (H1792) or 10nM (H23) G12Ci or 10nM G12Di (for A427, SKLU-1 and HCC461) or 10 µM AKTi and a combination of both in 3D. 48 hours later, viability was measured by CellTiter-Glo 3D. Viability expressed as % of DMSO control. Mean viability of each treatment response per cell line is depicted in the graph. CI values were calculated (n=3). (D) Cell death analyses of human KRAS^G12D^ NSCLC cell lines SKLU-1 and HCC461 in response to 10nM G12Di and 10µM AKTi and a combination of both in 3D. Cell death was measured after 48 hours of drug exposure by flow cytometric quantification of PI staining. Data normalised to DMSO control (n=3). Mean ±s.e.m. depicted for all graphs and statistical analysis carried out for C using two-way ANOVA and D using one-way ANOVA. ****P<0.0001, ***P<0.001, **P<0.01, *P<0.05, ns>0.05. Note: CI = combination index. For CI analysis, points appearing above the top dotted line signify drug antagonism. Points appearing between the top and bottom dotted line signify drug additivity. Points appearing below the bottom dotted line signify drug synergism.

We next exposed human H2030 (KRAS^G12C^) and HCC461 (KRAS^G12D^) NSCLC cell lines to increasing doses of G12Ci or G12Di respectively in the presence or absence of AKTi and again saw that the combination of G12Di + AKTi was synergistic across all doses (Figure 5B). The combination of G12Ci +AKTi was only synergistic at high doses of G12Ci, indicating that toxicity may preclude this combination as a clinical option (Figure 5B). We also exposed a different human KRAS^G12D^ cell line, SKLU-1, which is relatively more resistant to G12Di, to the same conditions. Similarly, we observed a synergistic response, highlighting a possible role for PI3K-AKT-mTOR axis disruption in overriding innate resistance to KRAS^G12D^ inhibition (Supplementary Figure S7C). Finally, we assessed our full panel of human KRAS^G12C^ and KRAS^G12D^ cell lines, exposing them to either a combination of G12Ci + AKTi or G12Di + AKTi. Using cell line-specific concentrations of either G12Ci or G12Di to elicit a comparable reduction in cell viability along with a single concentration of AKTi, the overall combined effect of G12Di and AKTi was significantly more potent than that of G12Ci and AKTi (Figure 5C). Moreover, the reduction in viability with the combination of G12Di and AKTi was due to apoptotic cell death which was further confirmed by rapid caspase-3/-7 activation (Figure 5D and Supplementary Figure S7D). In order to confirm that the synergism between G12Di and AKTi was not due to off-target effects, we exposed KRAS^G12C^ MEFs to this combination in which no further benefit was seen compared to AKTi alone (Supplementary Figure S7E). To further underpin the importance of inhibiting the PI3K-AKT-mTOR pathway to enhance KRAS^G12D^ inhibition, we co-inhibited KRAS^G12D^ and the MAPK pathway (ERKi), which resulted in a weakly additive response in KRAS^G12D^ cell lines (Supplementary Figure S7F). Altogether, these data show that rational selection of an up-front combination of KRAS mutant-specific inhibitors with inhibitors of mutant allele-specific vulnerabilities will achieve a greater therapeutic impact, minimising the risk of developing resistance. G12Di + AKTi conferred a strong therapeutic response relative to G12Di + ERKi in KRAS^G12D^ cell lines or G12Ci + AKTi in KRAS^G12C^ cell lines. Thus, in the context of KRAS^G12D^-driven NSCLC, we have identified a novel treatment vulnerability, patient selection strategy and combination approach.

## Discussion

KRAS mutant NSCLC heterogeneity limits treatment efficacy resulting in poor patient outcomes^3^. While it is becoming clear that KRAS mutations exhibit distinct biological properties^8,9,43^, determining mutant isoform-specific treatment vulnerabilities is under-researched. Here we show that KRAS^G12D^ is more potent than the more commonly occurring KRAS^G12C^ isoform at initiating lung tumorigenesis. We also demonstrate that this superior oncogenicity may be linked to hyperactive PI3K-AKT-mTOR signaling. Interestingly, we see that this initial difference in potency is lost through tumor progression and the immediate signaling differences diminish; however, certain signaling dependencies persist or emerge during progression offering therapeutically actionable targets (Figure 6). We propose that KRAS^G12C^ relies more on KRAS signaling through the MAPK arm to increase oncogenic potential and promote tumor growth rendering advanced KRAS^G12C^ tumors more susceptible to MAPK inhibition compared to KRAS^G12D^ tumors. In contrast, KRAS^G12D^ tumors rely on PI3K-AKT-mTOR signalling and are more vulnerable to inhibition of this pathway. Combination of KRAS^G12D^ and PI3K-AKT-mTOR pathway inhibition in KRAS^G12D^ cells was synergistic, eliciting a cytotoxic response that represents a potential mutant-specific treatment approach for NSCLC.

**Figure 6.**
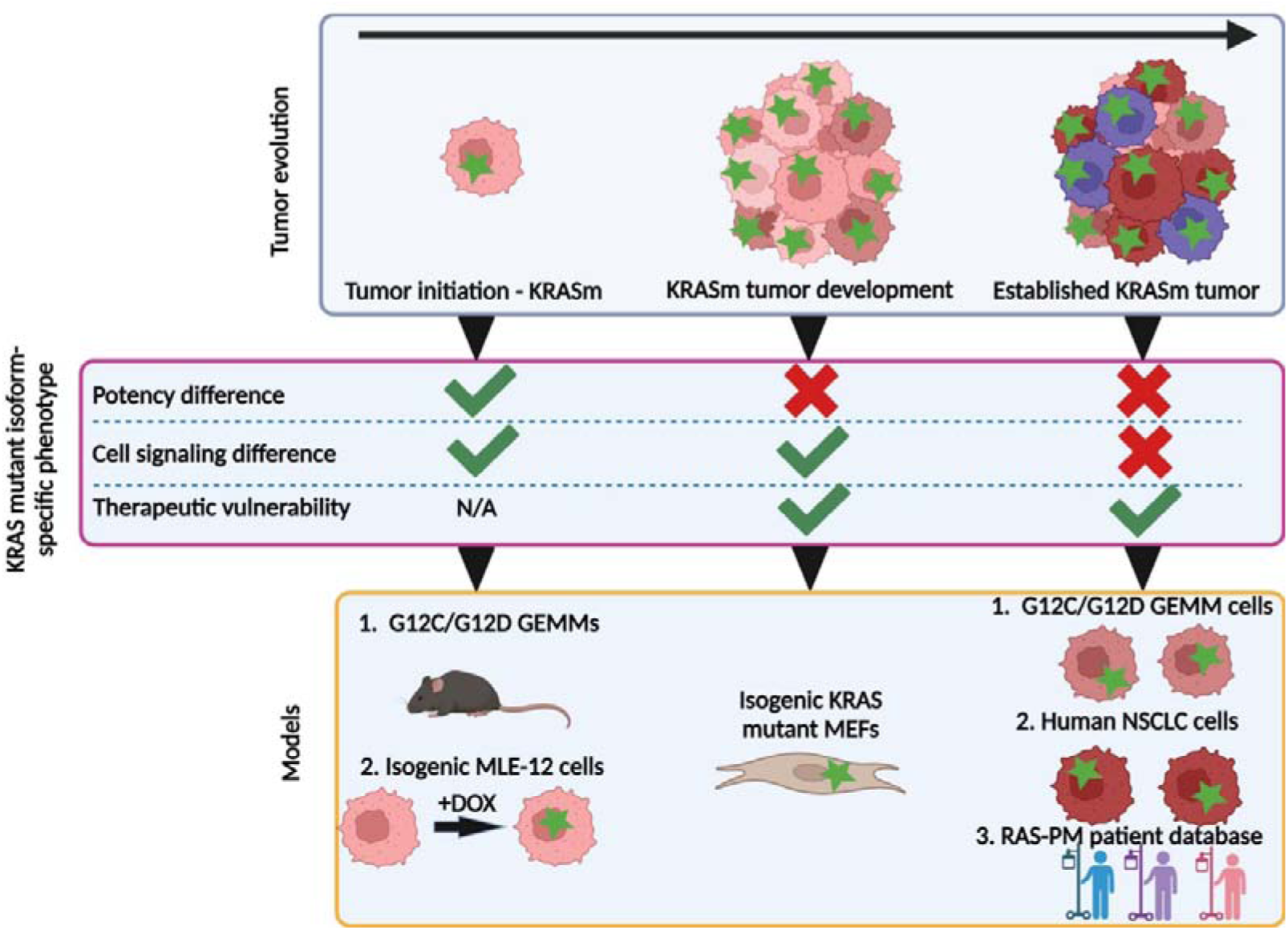
Schematic illustrating the progression of KRAS mutant-specific NSCLC. Upon initiation, the greater potency of KRAS^G12D^ induces rapid tumorigenesis relative to KRAS^G12C^ potentially via the PI3K-AKT-mTOR pathway. During progression, the KRAS mutant isoform-specific differences in potency are not evident, underpinning the equivalent survival outcomes in patients. However, KRAS^G12D^ tumors maintain reliance on the PI3K-AKT-mTOR pathway, while KRAS^G12C^ increases oncogenic potential through other means, conferring therapeutic vulnerabilities which can be exploited. Green star = KRAS mutation. Created with BioRender.com.

A striking finding from this study was the difference in lung tumor initiation and latency between mice harboring different KRAS-mutant alleles. Tumors of *Kras^G12D^* mice were more abundant and grew more rapidly, translating into poorer animal survival. Interestingly, these findings mirror those from a pancreatic cancer model in which at 12 weeks after KRAS mutant activation, *Kras^G12D^* mice showed more extensive pancreatic intraepithelial neoplasias (PanINs) compared to *Kras^G12C^* mice. In the same study, *Kras^G12C^* or *Kras^G12D^* activation in colonic epithelium had an equal tumorigenic response^44^. Collectively, these reported findings and our data support the concept that the oncopotencies of KRAS mutations are tissue-specific^9^, with *Kras^G12C^* and *Kras^G12D^* possessing contrasting potency in both the lung and pancreas, but similar potency in the colon. Our initiation model supported the *in vivo* phenotype, demonstrating that *Kras^G12D^* is more potent than *Kras^G12C^*at increasing proliferation. We also identified upregulated PI3K-AKT-mTOR signaling as a possible contributor to this more oncogenic phenotype associated with *Kras^G12D^*. Through inhibition of this pathway, we saw that the expression of markers associated with tumorigenesis was reduced to a greater extent in *Kras^G12D^* MLE-12 cells. Furthermore, *in vivo*, we saw higher S6 activation in *Kras^G12D^*-driven early lesions. In PDAC initiation, mTOR signaling was hyperactivated in *Kras^G12D^*-driven acinar-to-ductal metaplasia (ADM). Genetic ablation of mTOR signaling components abolished ADM initiation^45^. Together, this report and our data both highlight that *Kras^G12D^* may require input from PI3K-AKT-mTOR signaling to drive tumorigenesis compared to Kras^G12C^.

Importantly, our established tumor models and patient data reveal that in advanced disease, differences in KRAS mutant isoform-specific potency are no longer evident, mirroring findings from previous large cohort studies which failed to support KRAS-mutant allele-specific differences in outcomes^8,46,47^. KRAS mutant isoforms occur alongside distinct co-mutation patterns which ultimately affect signaling networks, immune surveillance and response to therapy^8,9^. Furthermore, *KRAS^G12C^* tumors have a higher mutational burden and are impacted by a higher number of co-mutations^8,9,20^. These genetic alterations are likely to compensate for the differences in mutant isoform-specific oncopotency. Thus, we propose that initially *KRAS^G12D^* is a stronger oncogene in NSCLC and promotes rapid tumor growth compared to *KRAS^G12C^*. However, over time, *KRAS^G12C^* tumors may acquire several additional alterations to increase tumorigenicity and, as a result, the difference in potency becomes less evident.

Our study also sheds light on the heterogeneity between KRAS mutant isoforms and isoform-specific RAS effector dependencies in established tumors. Both MEF and human NSCLC gene expression data showed enrichment of genes related to KRAS signaling (*KRAS^G12C^*) and PI3K-AKT-mTOR signaling (*KRAS^G12D^*). Interestingly, this did not align with phosphorylation levels of effector proteins within these pathways in our panel of NSCLC cell lines as they exhibited heterogeneous phosphorylation levels that were cell line-specific, rather than KRAS mutant isoform-specific. Surprisingly, despite the varied phosphorylation levels, we saw KRAS mutant isoform-specific responses to MAPK or PI3K-AKT-mTOR inhibition in NSCLC cell lines. We observed that ERK and MEK inhibition individually had a stronger impact on viability in *KRAS^G12C^* NSCLC lines compared to *KRAS^G12D^*. This is in agreement with a previous study using a MEK inhibitor in an isogenic MEF panel, which reported greater sensitivity of Kras^G12C^ MEFs to MEK inhibition despite exhibiting comparable levels of phosphorylated MEK with Kras^G12D^ MEFs^8^. Additionally, *KRAS^G12D^*NSCLC cell lines were more susceptible to PI3K-AKT-mTOR inhibition. The varied levels of PI3K-AKT-mTOR activation observed among NSCLC cell lines may be influenced by factors such as co-mutations. However, despite the heterogeneity in *KRAS^G12D^* cell lines, this pathway still acts as a critical node to maintain tumor viability. Clinical trials have returned disappointing results for agents targeting PI3K-AKT-mTOR signaling as monotherapy in NSCLC. However, these trials were carried out on molecularly unselected cohorts^48^. Despite AKT activating mutations being rare, AKT isoforms are overexpressed in NSCLC^49^ and there are reports of efficacy with AKT inhibition as part of combination treatments in lung cancer patients^50^, implying that there is potential for NSCLC patients to benefit from PI3K-AKT-mTOR targeted therapy. In light of this and our findings, we propose targeting PI3K-AKT-mTOR signaling in *KRAS^G12D^*-driven LUAD as a potential therapeutic option. Additionally, as our MEF and publicly available human CCLE gene expression data showed similar gene-set enrichment which informed treatment vulnerabilities, it is worth considering that genotype-specific transcriptional signatures are better determinants of therapeutic vulnerabilities rather than pathway activation.

As already mentioned, studies have shown that KRAS mutant isoforms exhibit distinct tissue-specific features. Thus, dissecting the individual functions of KRAS mutant isoforms in the context of NSCLC is critical to inform combination partners to maximise the effectiveness of direct KRAS inhibitors in this disease setting. There are emerging reports of effective combination treatments to increase the effectiveness of KRAS^G12D^ inhibition in colorectal cancer^41,42^. However, to our knowledge, there has not been any investigation into combinatorial strategies involving KRAS^G12D^ inhibition in NSCLC. We showed that the combination of KRAS^G12D^ inhibition and PI3K-AKT-mTOR pathway inhibition synergised in reducing cellular viability, whilst other combinations were mostly additive. Even more importantly, this synergy was not cell line-specific but was achieved in all the *KRAS^G12D^* cell lines we examined.

To conclude, our study emphasises that different KRAS mutations exhibit different oncopotencies which become less apparent over time despite retaining intrinsic dependencies resulting in therapeutic vulnerabilities. More specifically, it highlights that one amino acid difference between RAS point mutations dictates precision medicine approaches in NSCLC. Our data supports the idea that solely confirming the presence of KRAS mutation in a patient with NSCLC is insufficient to inform treatment options. Rather, knowing the type of KRAS mutation is critical especially now with the new wave of KRAS mutant isoform-specific inhibitors emerging which require combinatorial treatments to maximise therapeutic efficacy and decrease resistance.

## Acknowledgements

CRL is an IASLC Young Investigator and is supported by the Manchester Cancer Research Centre Town Hall Programme. This work was also supported by core funding to the CRUK Manchester Institute (CRUK MI) (grant A27412), Manchester CRUK Centre Award (grant A25254), CRUK Manchester Experimental Cancer Medicines Centre (grant A25146), CRUK Lung Cancer Centre of Excellence (grant A20465) and by The Christie Charitable Fund. We are grateful to the Genome Editing and Mouse Models facility of CRUK MI for the generation of conditional KRAS^G12C^ mice. We also thank all the other core facilities of CRUK MI for help with experiments especially the Biological Resources Unit and the Molecular Biology, Histology, Visualisation, Irradiation & Analysis and Flow Cytometry facilities. We thank Dominic Esposito’s lab at Frederick NCI for the constructs to generate the MLE-12 cell line panel and William Burgan also at Frederick NCI for supplying the KRAS mutant MEF panel.

## Author contributions

Conceptualization, W.J.M, L.W., C.R.L., A.M. (CRUK); methodology, W.J.M, L.W., J.S., A.M. (CRUK); validation, W.J.M, H.A.; formal analysis, W.J.M., M.J.B, J.S., T.B.; investigation, W.J.M, L.W., H.A., M.J.B. J.S, L.G., T.B.; resources, A.V., B.B., H.A.W., F.B., C.R.L., A.M. (CRUK); data curation, H.A., M.A., A.M. (GV), J.A., B.B., H.A.W.; writing–original draft, W.J.M., C.R.L., A.M. (CRUK); writing–review & editing, W.J.M, C.R.L., A.M. (CRUK); visualization, W.J.M., H.A., L.G., C.R.L., A.M. (CRUK); funding acquisition, F.B., C.R.L., A.M. (CRUK); supervision, C.R.L., A.M. (CRUK).

## References

1. Thomas, A., Liu, S. V., Subramaniam, D. S. & Giaccone, G. Refining the treatment of NSCLC according to histological and molecular subtypes. Nature Reviews Clinical Oncology vol. 12 511–526 (2015).

2. Siegel, R. L., Miller, K. D., Fuchs, H. E. & Jemal, A. Cancer statistics, 2022. CA Cancer J Clin 72, 7– 33 (2022).

3. Siegel, R. L., Miller, K. D. & Jemal, A. Cancer statistics, 2020. CA Cancer J Clin 70, 7–30 (2020).

4. Sainz de Aja, J., Dost, A. F. M. & Kim, C. F. Alveolar progenitor cells and the origin of lung cancer. J Intern Med 289, 629–635 (2021).

5. Salgia, R., Pharaon, R., Mambetsariev, I., Nam, A. & Sattler, M. The improbable targeted therapy: KRAS as an emerging target in non-small cell lung cancer (NSCLC). Cell Reports Medicine vol. 2 (2021).

6. Collins, M. A. et al. Oncogenic Kras is required for both the initiation and maintenance of pancreatic cancer in mice. Journal of Clinical Investigation 122, 639–653 (2012).

7. Judd, J. et al. Characterization of KRAS mutation subtypes in non-small cell lung cancer. Mol Cancer Ther 20, 2577–2584 (2021).

8. Ricciuti, B. et al. Comparative Analysis and Isoform-Specific Therapeutic Vulnerabilities of KRAS Mutations in Non-Small Cell Lung Cancer. Clinical Cancer Research 28, 1640–1650 (2022).

9. Cook, J. H., Melloni, G. E. M., Gulhan, D. C., Park, P. J. & Haigis, K. M. The origins and genetic interactions of KRAS mutations are allele- and tissue-specific. Nat Commun 12, (2021).

10. Miller, M. S. & Miller, L. D. RAS mutations and oncogenesis: Not all RAS mutations are created equally. Frontiers in Genetics vol. 2 (2012).

11. Li, S., Balmain, A. & Counter, C. M. A model for RAS mutation patterns in cancers: finding the sweet spot. Nature Reviews Cancer vol. 18 767–777 (2018).

12. Jänne, P. A. et al. Selumetinib plus docetaxel compared with docetaxel alone and progression-free survival in patients with KRAS-mutant advanced non-small cell lung cancer: The SELECT-1 randomized clinical trial. JAMA - Journal of the American Medical Association 317, 1844–1853 (2017).

13. Ostrem, J. M., Peters, U., Sos, M. L., Wells, J. A. & Shokat, K. M. K-Ras(G12C) inhibitors allosterically control GTP affinity and effector interactions. Nature 503, 548–551 (2013).

14. Canon, J. et al. The clinical KRAS(G12C) inhibitor AMG 510 drives anti-tumour immunity. Nature 575, 217–223 (2019).

15. Skoulidis, F. et al. Sotorasib for Lung Cancers with KRAS p.G12C Mutation . New England Journal of Medicine 384, 2371–2381 (2021).

16. Riely, G. J. et al. 99O PR KRYSTAL-1: Activity and preliminary pharmacodynamic (PD) analysis of adagrasib (MRTX849) in patients (Pts) with advanced non–small cell lung cancer (NSCLC) harboring KRASG12C mutation. Journal of Thoracic Oncology 16, S751–S752 (2021).

17. Wang, X. et al. Identification of MRTX1133, a Noncovalent, Potent, and Selective KRASG12DInhibitor. J Med Chem 65, 3123–3133 (2022).

18. Dunnett-Kane, V., Nicola, P., Blackhall, F. & Lindsay, C. Mechanisms of resistance to krasg12c inhibitors. Cancers vol. 13 1–14 (2021).

19. Liu, C. et al. KRAS-G12D mutation drives immune suppression and the primary resistance of anti-PD-1/PD-L1 immunotherapy in non-small cell lung cancer. Cancer Commun 42, 828–847 (2022).

20. Gao, G. et al. KRAS G12D mutation predicts lower TMB and drives immune suppression in lung adenocarcinoma. Lung Cancer 149, 41–45 (2020).

21. Ricciuti, B. et al. Dissecting the clinicopathologic, genomic, and immunophenotypic correlates of KRASG12D-mutated non-small-cell lung cancer. Annals of Oncology 33, 1029–1040 (2022).

22. Richardson, C. D., Ray, G. J., DeWitt, M. A., Curie, G. L. & Corn, J. E. Enhancing homology-directed genome editing by catalytically active and inactive CRISPR-Cas9 using asymmetric donor DNA. Nat Biotechnol 34, 339–344 (2016).

23. Jackson, E. L. et al. Analysis of lung tumor initiation and progression using conditional expression of oncogenic K-ras. Genes Dev 15, 3243–3248 (2001).

24. Jackson, E. L. et al. The differential effects of mutant p53 alleles on advanced murine lung cancer. Cancer Res 65, 10280–10288 (2005).

25. Wikenheiser, K. A., et al. Production of immortalized distal respiratory epithelial cell lines from surfactant protein C/simian virus 40 large tumor antigen transgenic mice. Proc. Natl. Acad. Sci. USA vol. 90 (1993).

26. Barbosa, M. A. G., Xavier, C. P. R., Pereira, R. F., Petrikaitė, V. & Vasconcelos, M. H. 3D Cell Culture Models as Recapitulators of the Tumor Microenvironment for the Screening of Anti-Cancer Drugs. Cancers vol. 14 (2022).

27. Sanaei, M. J., Razi, S., Pourbagheri-Sigaroodi, A. & Bashash, D. The PI3K/Akt/mTOR pathway in lung cancer; oncogenic alterations, therapeutic opportunities, challenges, and a glance at the application of nanoparticles. Translational Oncology vol. 18 (2022).

28. Zhao, P. et al. CD44 promotes Kras-dependent lung adenocarcinoma. Oncogene 32, 5186–5190 (2013).

29. Hase, T. et al. Pivotal role of epithelial cell adhesion molecule in the survival of lung cancer cells. Cancer Sci 102, 1493–1500 (2011).

30. Cui, Y. et al. Dynamic Expression of EpCAM in Primary and Metastatic Lung Cancer Is Controlled by Both Genetic and Epigenetic Mechanisms. Cancers (Basel) 14, (2022).

31. Drosten, M. et al. Genetic analysis of Ras signalling pathways in cell proliferation, migration and survival. EMBO Journal 29, 1091–1104 (2010).

32. Waters, A. M. et al. Evaluation of the selectivity and sensitivity of isoform-And mutation-specific RAS antibodies. Sci Signal 10, (2017).

33. Chung, W. J. et al. Kras mutant genetically engineered mouse models of human cancers are genomically heterogeneous. Proc Natl Acad Sci U S A 114, E10947–E10955 (2017).

34. Gao, J. et al. Integrative analysis of complex cancer genomics and clinical profiles using the cBioPortal. Sci Signal 6, (2013).

35. Cerami, E. et al. The cBio Cancer Genomics Portal: An open platform for exploring multidimensional cancer genomics data. Cancer Discov 2, 401–404 (2012).

36. Jamal-Hanjani, M. et al. Tracking the Evolution of Non–Small-Cell Lung Cancer. New England Journal of Medicine 376, 2109–2121 (2017).

37. Campbell, J. D. et al. Distinct patterns of somatic genome alterations in lung adenocarcinomas and squamous cell carcinomas. Nat Genet 48, 607–616 (2016).

38. Dongre, A. & Weinberg, R. A. New insights into the mechanisms of epithelial–mesenchymal transition and implications for cancer. Nature Reviews Molecular Cell Biology vol. 20 69–84 (2019).

39. Barretina, J. et al. The Cancer Cell Line Encyclopedia enables predictive modelling of anticancer drug sensitivity. Nature 483, 603–607 (2012).

40. Zhang, J. et al. Resistance looms for KRAS G12C inhibitors and rational tackling strategies. Pharmacology and Therapeutics vol. 229 (2022).

41. Hallin, J. et al. Anti-tumor efficacy of a potent and selective non-covalent KRASG12D inhibitor. Nat Med 28, 2171–2182 (2022).

42. Feng, J. et al. Feedback activation of EGFR/wild-type RAS signaling axis limits KRASG12D inhibitor efficacy in KRAS G12D-mutated colorectal cancer. Oncogene (2023).

43. Muñoz-Maldonado, C., Zimmer, Y. & Medová, M. A comparative analysis of individual ras mutations in cancer biology. Front Oncol 9, (2019).

44. Zafra, M. P. et al. An in vivo kras allelic series reveals distinct phenotypes of common oncogenic variants. Cancer Discov 10, 1654–1671 (2020).

45. Zhao, Y. et al. mTORC1 and mTORC2 Converge on the Arp2/3 Complex to Promote KrasG12D-Induced Acinar-to-Ductal Metaplasia and Early Pancreatic Carcinogenesis. Gastroenterology 160, 1755–1770.e17 (2021).

46. Arbour, K. C. et al. Treatment outcomes and clinical characteristics of patients with KRAS-G12C-mutant non-small cell lung cancer. Clinical Cancer Research 27, 2209–2215 (2021).

47. Bironzo, P. et al. Real world retrospective study of KRAS mutations in advanced non–small cell lung cancer in the era of immunotherapy. Cancer 129, 1662–1671 (2023).

48. Tan, A. C. Targeting the PI3K/Akt/mTOR pathway in non-small cell lung cancer (NSCLC). Thoracic Cancer vol. 11 511–518 (2020).

49. Scrima, M. et al. Signaling networks associated with AKT activation in non-small cell lung cancer (NSCLC): New insights on the role of phosphatydil-inositol-3 kinase. PLoS One 7, (2012).

50. Lara, P. N. et al. Phase II study of the AKT inhibitor MK-2206 plus erlotinib in patients with advanced non-small cell lung cancer who previously progressed on erlotinib. Clinical Cancer Research 21, 4321–4326 (2015).

