## Supplementary file for "The PI3K-AKT-mTOR axis persists as a therapeutic dependency in KRAS^G12D^-driven non-small cell lung cancer"

**Supplementary materials and methods**

**Generation of *Kras^G12C^* mouse model.** To generate a mouse model carrying the conditional expression of Kras^G12C^, we employed CRISPR-Cas9 technology to the *Kras^LSL-G12D^* mouse model from Tyler Jacks (JAX stock #008179). Briefly, in order to design specific guide RNAs and oligos, the region around *Kras^LSL-G12D^* point mutation was fully sequenced. Using the CRISPR gRNA tool from MIT, a gRNA was designed that targets the wild type version of Kras but modified with the nucleotide mutation present in the *Kras^LSL-G12D^* model (Supplementary Table S1). Complementary oligos carrying the specific 20-nucleotide sequence of the gRNA were annealed and cloned into a vector containing the gRNA backbone and a T7 promoter for *in vitro* transcription using MEGAshortscript T7 Kit (Life Technologies, #AM1354) and purified with MEGAclear Kit (Life Technologies, #AM1908). The repair template was designed as an Ultramer (IDT DNA technologies) single stranded, asymmetric DNA donor, complementary to the strand that the gRNA binds to following a previously described method^22^. To facilitate screening of targeted events by PCR-RFLP, HaeIII restriction silent mutation was inserted. Superovulated female C57BL/6OlaHsd mice (4-weeks old) were mated with C57BL/6OlaHsd stud males, and fertilized embryos were collected from oviducts. Mice were generated by cytoplasmic microinjection of 100ng/µl mRNA Cas9 (TriLink, #L-7206), 25ng/µl *in vitro* transcribed gRNA and Ultramer ssDNA (100ng/µl) repair template into 1-cell stage C57BL/6OlaHsd embryos. Microinjected embryos were surgically implanted into the oviduct of day 0.5 post coitum Hsd:ICR (CD-1^®^) pseudopregnant mice. The offspring was assessed for correct targeting using PCR-RFLP primers flanking the homology arms (Supplementary Table S1) followed by HaeIII restriction digestion (New England Biolabs) and subsequent Sanger sequencing. 3 mice carrying the correct targeting mutation were selected and founder lines were established (KRC, KRCA and KRCB). Germline transmission for each founder was confirmed using the same approach and independent colonies were established. New model now termed Kras^LSL-G12C^ mouse model.

***In vivo* studies.** Kras^LSL-G12C^ and Kras^LSL-G12D^ mice were cross-bred with t*p53^KO^* mice to obtain Kras^LSL-G12C^/tp53^fx/fx^ and Kras^LSL-G12D^/tp53^fx/fx^. Endogenous lung tumours were generated through intranasal or intratracheal administration of 8–16-week-old Kras^LSL-G12C^/tp53^fx/fx^ and Kras^LSL-G12D^/tp53^fx/fx^ with adenovirus expressing Cre recombinase (AdVC; University of Iowa Vector Core, USA) establishing Kras^G12C^/tp53^KO^ and Kras^G12D^/tp53^KO^ models.

For the tumor burden study to confirm the Kras^G12C^ mouse model worked, mice were culled 1 month after instillation of AdVC at 1x10^8^ plaque forming units (PFU) per mouse. For comparative tumor burden studies, mice were culled either 11 months (for t*p53^WT^* mice) or 4 months (for *tp53^KO^* mice) after AdVC inhalation (1 × 10^6^ PFU per mouse). For the survival study, mice were instilled with 1 × 10^7^ PFU per mouse and were culled upon onset of moderate symptoms of disease. For the early lesions study, mice were instilled with 1 × 10^7^ PFU per mouse and were culled between 2 and 6 weeks post instillation. Lungs were collected at study endpoints and formalin fixed for 24-hours, following which lungs were stored in ethanol for histological analyses. Mice were randomised with respect to age and gender.

**Histological analyses.** Fixed lung samples were embedded in paraffin blocks (FFPE). Lung sections were stained with haematoxylin and eosin (H&E) using standard methods. Whole slide imaging was performed using an Olympus VS200 MTL (Olympus Tokyo, Japan). Quantification of lung tumors and hyperplasia was performed using HALO® software (Indica Labs, New Mexico USA). Lung tissue was classified to differentiate tumor and hyperplastic tissue from other tissue. Tumor and hyperplasia lung areas were calculated with the following formula: tumour area (or hyperplastic area)/ total lung area × 100%. For calculation of tumor number, tumors with length greater than 200µm in diameter were included. For staining of murine lung tumors from GEMM models by immunohistochemistry (IHC), FFPE samples were cut as 4µm sections and stained for markers using antibodies at optimised dilutions detailed in Supplementary Table S2. All IHC was carried out using the BOND RX automated platform (Leica Microsystems). Sections were mounted on charged slides, followed by de-paraffinization. Heat-induced epitope retrieval was accomplished using Epitope Retrieval Solution 1 (Leica Microsystems, AR9961) for 20-minutes at 98°C. Using the BOND Polymer Refine Detection kit (Leica Microsystems, DS9800), endogenous peroxidase was blocked using 3% v/v hydrogen peroxide for 10-minutes and the slides were further blocked with 10% w/v casein (Vector, SP5020) in TBS/T. Antibodies were applied for 30-minutes, visualised and then counterstained with haematoxylin. Whole slide imaging was performed using the Olympus VS200 MTL and chromogenic staining was quantified using HALO® software. Immunostaining within tumors and early lesions was based on matched serial H&E sections. The immunostaining of total protein (Cyclin D1 and Ki67) within tumors was quantified as % positive, while immunostaining of phosphorylated proteins (p-S6 and p-ERK) were quantified as a gradient of low, medium and high % positive, with high % positive reported. For the early lesions study, samples were excluded from analyses if less than 20 tumors were counted on corresponding H&E section.

**Generation of isogenic KRAS mutant MLE-12 cell panel.** Gateway recombination cloning was used to assemble multiple entry clones into a final destination vector. Entry clones used were: AttL4-AttR5 tetracycline inducible (TRE3G) promoter, AttL5-AttR1 N-terminal FLAG tag and AttL1-AttL2 human KRAS4b constructs (*KRAS^WT^*, *KRAS^G12C^* or *KRAS^G12D^*). The destination vector used was AttR4-AttR2 DEST362 lentiviral mammalian expression vector. Gateway LR clonase (ThermoFisher, #11791019) reaction was performed as per manufacturer’s protocol using 50ng entry clones and 100ng destination vector at 25°C overnight, then inactivated with proteinase K at 37°C for 15 minutes. Following bacterial transformation of reaction mixes in DH10B competent cells, colonies were selected, and gateway recombination confirmed by Sanger sequencing. Lentivirus was produced for each construct using Lenti-X 293T cells and second generation lentiviral packaging plasmids. MLE-12 cells were transduced with lentivirus and 10μg/ml polybrene and successfully transduced cells were selected with 5μg/ml puromycin.

**Generation of GEMM-derived cell lines.** Lungs with tumors from *Kras^G12C^/tp53^KO^* and *Kras^G12D^/tp53^KO^* mice were removed and cut into small pieces and cultured in appropriate media. Immune cells and fibroblasts died over the course of the initial passages. A pure population was confirmed by flow cytometric staining of pan-cytokeratin.

**Reagents.** Sotorasib (KRAS^G12C^ inhibitor, #HY-114277), Ulixertinib (ERK 1/2 inhibitor, #HY-15816), U0126 (MEK1 and MEK2 inhibitor, #HY-12031A), Buparlisib (pan-class I PI3K inhibitor, #HY70063), Rapamycin (mTORC1 inhibitor, #HY-10219) and Everolimus (mTOR inhibitor, #HY-10218) were all purchased from MedchemExpress. MRTX1133 (KRAS^G12D^ inhibitor, #CAY-36413) was purchased from Cayman Chemicals. Capivasertib (AKT inhibitor, #S8019) was purchased from Selleck Chemicals. Doxycycline (#D9891) was purchased from Sigma. 96-well ultra-low attachment (ULA) plates (#7007), 24-well ULA plates (#3473) and 6-well ULA plates (#3471) were purchased from Corning.

**Proliferation and viability assays.** For MEFs, proliferation was measured by three methods – (1) using the Cell Transformation kit (Abcam, #Ab235698) as per manufacturer’s guidelines; (2) by measuring spheroid area under ultra-low attachment (ULA) conditions in 96-well ULA plates using ImageJ software; and (3) by labelling cells with 1 µM BrdU (Sigma, #B5002) for 6 hrs in 6-well ULA plates, followed by fixation in ethanol and staining with FITC-conjugated anti-BrdU monoclonal antibody (Biolegend, #3641034) and propidium iodide (PI). BrdU-positive cells were measured using a BD Fortessa flow cytometer and analysed with FlowJo software (Treestar, USA) and expressed as % of total cell population. For drug responses, MEFs were seeded under ULA conditions in 96-well ULA plates and after indicated time points, viability was measured using CellTiter-Glo® 3D (Promega, cat# G9681) according to manufacturer’s guidelines. For GEMM-derived cell lines (mTCLs), proliferation and viability in response to drug treatment were measured under attached conditions and, after the indicated timepoints, cells were fixed with crystal violet and the absorbance of the retained dye was measured at 595 nm. For MLE-12 cells, proliferation and viability in response to doxycycline treatment were measured by two methods – (1) by measuring spheroid area under ULA conditions in 96-well ULA plates using ImageJ software; and (2) by growing under ULA conditions in 96-well ULA plates and using CellTiter-Glo® 3D according to manufacturer’s guidelines. For human NSCLC cell lines, proliferation and viability in response to drug treatment was measured under ULA conditions in 96-well ULA plates and by using CellTiter-Glo® 3D according to manufacturer’s guidelines. Unless stated otherwise, drug treatments were carried out 24 hours after seeding.

**Combination indices.** Combination indices were calculated using the coefficient of drug interaction (CDI) equation which is defined as: CDI=AB/(AxB), where A is the relative viability of cells after exposure to drug A, B is the relative viability of cells after exposure to drug B and AB is the relative viability of cells after exposure to drug A and drug B in combination. Relative viability is expressed as a ratio of drug treated cells compared to the DMSO control. CDI value of >1.1 denotes drug antagonism; 0.9-1 denotes additivity; and <0.9 denotes synergism.

**Western blotting.** Cells were seeded under ULA conditions using 6-well ULA plates and after the indicated time point, cells were lysed in RIPA buffer (Sigma, #R0278) containing PhosSTOP (Roche, #4906845001) and protease inhibitor cocktail (Sigma, #P8340)and protein lysates were collected, resolved by SDS-PAGE using 4%–12% Bis-Tris NuPAGE gels in MOPs running buffer and transferred to PVDF membranes (Merck, #IPVH00010). Membranes containing the separated proteins were blocked in 5% bovine serum albumin dissolved in phosphate-buffered saline and tween (BSA-PBS/T) and then incubated with primary antibodies overnight. Membranes were then incubated with horse radish peroxidase-conjugated secondary antibodies to enable target protein detection using ECL substrate (Pierce, #32106) and a BioRad Chemidoc imager. Primary and secondary antibodies used for Western blotting are outlined in Supplementary Table S2.

**RNA-sequencing and analyses.** Cells were seeding under ULA conditions in 6-well ULA plates and after the indicated time point, RNA was extracted from cells using the RNeasy Kit (Qiagen, #74104) following the instructions of the manufacturer. SureSelect (Agilent; full-length RNA; paired-end) cDNA libraries were prepared and sequenced using the Novaseq platform (Illumina). Basecalls were converted to fastq files using bcl2fastq (Illumina). Fastq files were aligned to GRCm38 (Ensembl75) using Star aligner (version 2.5.1b). BAM alignments were quantified in R (version 3.6.1) using featureCounts from the Rsubread library (version 2.0.1). QC of the count data was assessed via PCA. DE analysis was performed using DESeq2 (version 1.26). Pre-ranked geneset enrichment analysis (GSEA) using the hallmark signature database (msigdbr; version 7.5.1) was performed with the fgsea package (version 1.24). For pairwise comparisons, the ranking metric for DEGs (pAdj<0.05) was calculated as [-log10(pAdj) x sign(log2FC)]; where a geneset was the product of two or more comparisons, consensus DEGs were screened for consistent directionality between comparisons and a directional p rank sum calculated as the ranking metric. Heatmaps of z-scores with hierarchical clustering were generated from TPM values of genes considered to be differentially expressed between conditions (log2-fold change ≥ 0.5 or ≤ -0.5, p ≤ 0.05). Gene sets from GSEA were used to define the genes included in each heatmap as indicated.

RNA-sequencing data of human KRAS^G12C^ and KRAS^G12D^ NSCLC cell lines was downloaded from Cancer Cell Line Encyclopedia (CCLE; <https://portals.broadinstitute.org/ccle>). Differential expression analysis was performed using the DESeq2 package (version 1.22.2) in R (version 3.5.1). For pathway enrichment analysis, genes that were significantly differentially expressed between KRAS^G12C^ and KRAS^G12D^ cell lines (false discovery rates (FDR) p value < 0.1) were compared against the Hallmark Database (Msigdb, Broad Institute, <https://www.gsea-msigdb.org/gsea/msigdb/index.jsp>). KRAS^G12C^ cell lines found in the database were: H358, H23, H2122, HCC1171, HCC44, SW1573, H2023, H1792, HOP62 and H1373. KRAS^G12D^ cell lines found in the database were: A427, SKLU-1 and HCC461.

**Intracellular staining.** Cells were seeded under ULA conditions in 6-well ULA plates (for MLE-12 cells, 100 ng/mL doxycycline was added at the time of seeding). 24-hours later, cells were harvested using PBS/ 0.5% EDTA and were fixed in paraformaldehyde. Cells were permeabilised using Triton-X and were stained with the indicated primary and fluorophore-conjugated secondary antibodies (Supplementary Table S2). Fluorescence intensity of fluorophore was measured using BD Fortessa flow cytometer recording 10,000 events per condition. Data was analysed using FlowJo10.

**Cell surface staining.** MLE-12 cells were seeded under ULA conditions in 6-well ULA plates and were exposed to 100 ng/mL doxycycline with/without either 1µM ERKi or 1 µM AKTi. After 24-hours, cells were harvested using PBS/ 0.5% EDTA and were stained with indicated fluorophore-conjugated antibodies and isotype control antibodies (Supplementary Table S2) in PBS/2.5% FBS. Both percentage positivity of fluorophore by gating on isotype control and median fluorescence intensity of fluorophore were measured using BD Fortessa flow cytometer recording 10,000 events per condition. Data was analysed using FlowJo10.

**Caspase-3/-7 detection.** Cells were seeded under ULA conditions in 96-well ULA plates and were treated 24 hours later with indicated compounds. 6 hours later, caspase-3/7 activation was measured by Caspase-Glo®3/7 (Promega, #G8090).

**Propidium iodide staining.** Cells were seeded under ULA conditions in 24 well ULA plates. 24-hours later, cells were treated with indicated compounds. 48 hours later, cells were harvested and stained with propidium iodide. Cells were analysed on BD Fortessa and cell death was calculated by gating on PI-positive cells and expressing PI-positive cells as a % of total cell population. Relative cell death was then further calculated by dividing the % cell death of compound treated cells by that of DMSO treated cells.

**cBioPortal exploration**

Publicly available data on cBioPortal was explored to evaluate tumor stage (T stage) as per the American Joint Committee on Cancer. Groups were stratified via KRAS G12C and KRAS G12D mutations.

**Supplementary figures**

**Supplementary Figure S1**

(A) (Left) Representative H&E section and (Right) HALO quantification of lung tumor area and hyperplasia for *Kras^G12C^/tp53^KO^* mice 4 months after exposure to AdVC (n=3 mice); scale bar = 5mm.
(B) HALO quantification of hyperplasia in lungs of *Kras^G12C^/tp53^WT^* and *Kras^G12D^/tp53^WT^* mice 11 months after exposure to AdVC (n=10 mice for *Kras^G12C^/tp53^WT^* genotype and 5 mice for *Kras^G12D^/tp53^WT^* genotype).
(C) HALO quantification of hyperplasia in lungs of *Kras^G12C^/tp53^KO^* and *Kras^G12D^/tp53^KO^* mice 4 months after exposure to AdVC (n=5 mice per genotype).
(D) (Left) Survival analysis and (Right) representative H&E sections of *Kras^G12C^/tp53^KO^* and *Kras^G12D^/tp53^KO^* mouse lungs after intratracheal delivery of AdVC (n=5 mice per genotype, Log-Rank (Mantel-Cox); scale bar = 5mm.
(E) HALO quantification of hyperplasia in lungs of *Kras^G12C^/tp53^KO^* and *Kras^G12D^/tp53^KO^* mice from survival study (n=5 mice per genotype).
(F) Mean tumor diameter per mouse per genotype taken from survival study (Figure 1E). Numbers above indicate overall mean per genotype (n=5 per genotype).

Mean ±s.e.m. depicted for A, B, C and E and statistical analysis (for B, C and E) carried out using using unpaired student’s t-test. ***P<0.001, **P<0.01, ns>0.05

**Supplementary Figure S2**

(A) Western blot analysis of KRAS, KRAS^G12D^ and ERK phosphorylation levels upon 24-hour exposure of isogenic MLE-12 cells to 100ng/mL doxycycline and 4-hour exposure to 1µM G12Ci.
(B) Representative images of MLE-12 spheroids upon treatment with 100ng/mL doxycycline for 96-hours. Scale bar = 400µm.
(C) Heatmaps showing DEGs belonging to mTORC1 Signaling gene set comparing each isogenic MLE-12 cell line after exposure to 100ng/mL doxycycline to its respective untreated control.
(D) Heatmaps showing DEGs belonging to Biocarta MAPK gene set comparing *KRAS^G12C^* to *KRAS^WT^* MLE-12 cells or *KRAS^G12D^* to *KRAS^WT^* MLE-12 cells 24 hours after exposure to 100ng/mL doxycycline.
(E) Western blot analysis of ERK and AKT phosphorylation levels upon 24-hour exposure of parental MLE-12 cells to 100ng/mL doxycycline. Representative of two independent experiments.

**Supplementary Figure S3**

(A) (Left) Representative images and (Right) quantification of viability of isogenic MEFs plated in agarose. Data normalised to day 0 readings (n=3). Scale bar = 1000µm.
(B) Representative images of spheroids of isogenic MEFs over time. Scale bar = 400µm.
(C) Flow cytometric quantification of S6 phosphorylation levels in isogenic MEFs 24-hours after seeding in 3D. Data normalised to *Kras^WT^* MEF (n=3).

A and C mean ±s.e.m. depicted and statistical analysis carried out using one-way ANOVA. **P<0.01, *P<0.05, ns>0.05

**Supplementary Figure S4**

Growth curves of *Kras^G12C^* and *Kras^G12D^* mTCL measured by crystal violet staining (n=3). Graph depicts mean ±s.e.m. and statistical analysis carried out using unpaired student’s t-test. ns>0.05

**Supplementary Figure S5**

(A) Schematic illustrating the proportion of patients with specific KRAS mutations enrolled in the RAS-PM database.
(B) 1^st^ line progression-free survival (PFS) of advanced NSCLC patients based on KRAS^G12C/G12D^ status. Log-Rank (Mantel-Cox) test was used to determine significance.
(C) Overall survival (OS) of advanced NSCLC patients based on KRAS^G12C/G12D^ status. Log-Rank (Mantel-Cox) test was used to determine significance.
(D) Schematic detailing which studies from cBioPortal examined for Tumour (T) stage evaluation.
(E) Tumour stage (T1-T4) of KRAS^G12C^ and KRAS^G12D^ NSCLC samples from cBioPortal. KRAS^G12C^: T1 56/139 (40.3%), T2 70/139 (50.4%), T3 9/139 (6.5%), T4 4/139 (2.9%). KRAS^G12D^: T1 19/35 (54.3%), T2 9/35 (25.7%), T3 4/35 (11.4%), T4 3/35 (8.6%).

**Supplementary Figure S6**

(A) Viability of human KRAS^G12C^ and KRAS^G12D^ NSCLC lines 72-hours after seeding in 3D measured using CellTiter-Glo 3D. Data normalised to HCC1171 (n=3).
(B) Significantly enriched Hallmark gene sets comparing *KRAS^G12C^* to *KRAS^G12D^* human NSCLC cell lines. Gene expression data downloaded from CCLE database. Arrows indicate gene sets of interest.
(C) Viability of human *KRAS^G12C^* and *KRAS^G12D^* NSCLC cell lines in response to 10µM MEKi in 3D. Viability was measured after 72-hours of drug exposure by CellTiter-Glo 3D (n=3).

A and C mean ±s.e.m. depicted and statistical analysis carried out using unpaired student’s t-test. *P<0.05, ns>0.05

**Supplementary Figure S7**

(A) A concentration point from each condition taken from Figure 5A per MEF line and plotted as bar chart.
(B) KRAS mTCLs were treated with either 10nM G12Ci (for *Kras^G12C^* mTCL), 1µM G12Di (for *Kras^G12D^* mTCL) and 20µM AKTi and a combination of both in 3D. 48 hours later, viability was measured by CellTiter-Glo 3D. Viability expressed as % of DMSO control. CI values were calculated (n=3).
(C) SKLU-1 (KRAS^G12D^) NSCLC cell line was treated with increasing concentrations of G12Di in the presence or absence of 10µM AKTi in 3D. 48 hours later, viability was measured by CellTiter-Glo 3D. Viability expressed as % of DMSO control. CI values were calculated (n=3).
(D) Caspase 3/7 activity analyses of human KRAS^G12D^ NSCLC cell lines SKLU-1 and HCC461 in response to 10nM G12Di and 10µM AKTi and a combination of both in 3D. Caspase 3/7 activity was measured after 6-hours of drug exposure using Caspase-Glo® 3/7 assay. Data normalised to DMSO control (n=3).
(E) KRAS^G12C^ MEFs were treated with either 1nM G12Di or 10µM AKTi and a combination of both in 3D. 48 hours later, viability was measured by CellTiter-Glo 3D. Viability expressed as % of DMSO control (n=3).
(F) Human KRAS^G12D^ NSCLCs (A427, SKLU-1 and HCC461) were treated with 10nM G12Di, 10µM ERKi and a combination of both in 3D. 48 hours later, viability was measured by CellTiter-Glo 3D. Viability expressed as % of DMSO control CI values were calculated (n=3).

Mean ±s.e.m. depicted for all graphs and statistical analysis carried out for A and B using two-way ANOVA and D, E and F using one-way ANOVA. ****P<0.0001, ***P<0.001, **P<0.01, *P<0.05, ns>0.05.

Note: CI = combination index. For CI analysis, points appearing above the top dotted line signify an antagonistic drug effect. Points appearing between the top and bottom dotted line signify an additive drug effect. Points appearing below the bottom dotted line signify a synergistic drug effect.

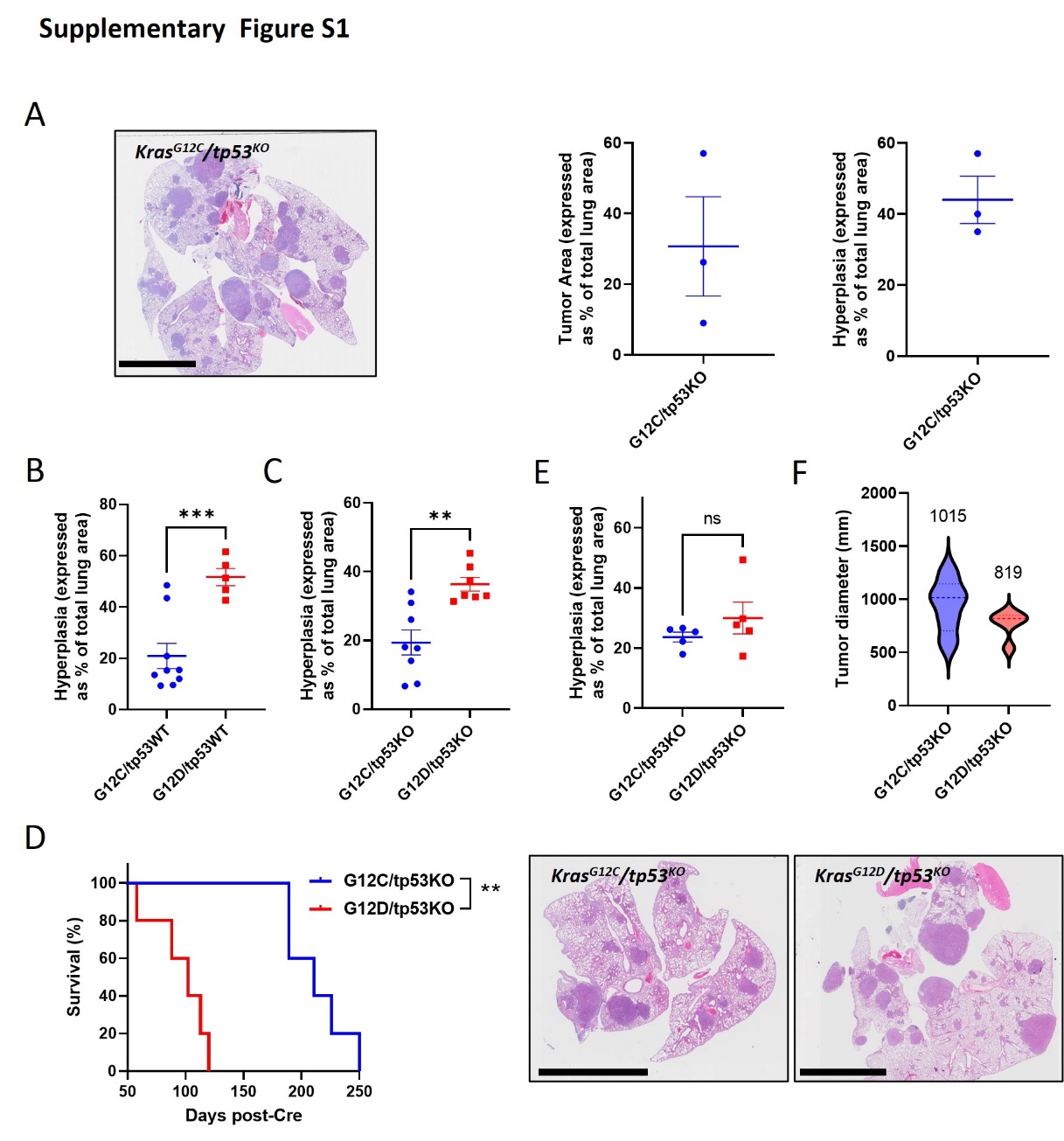

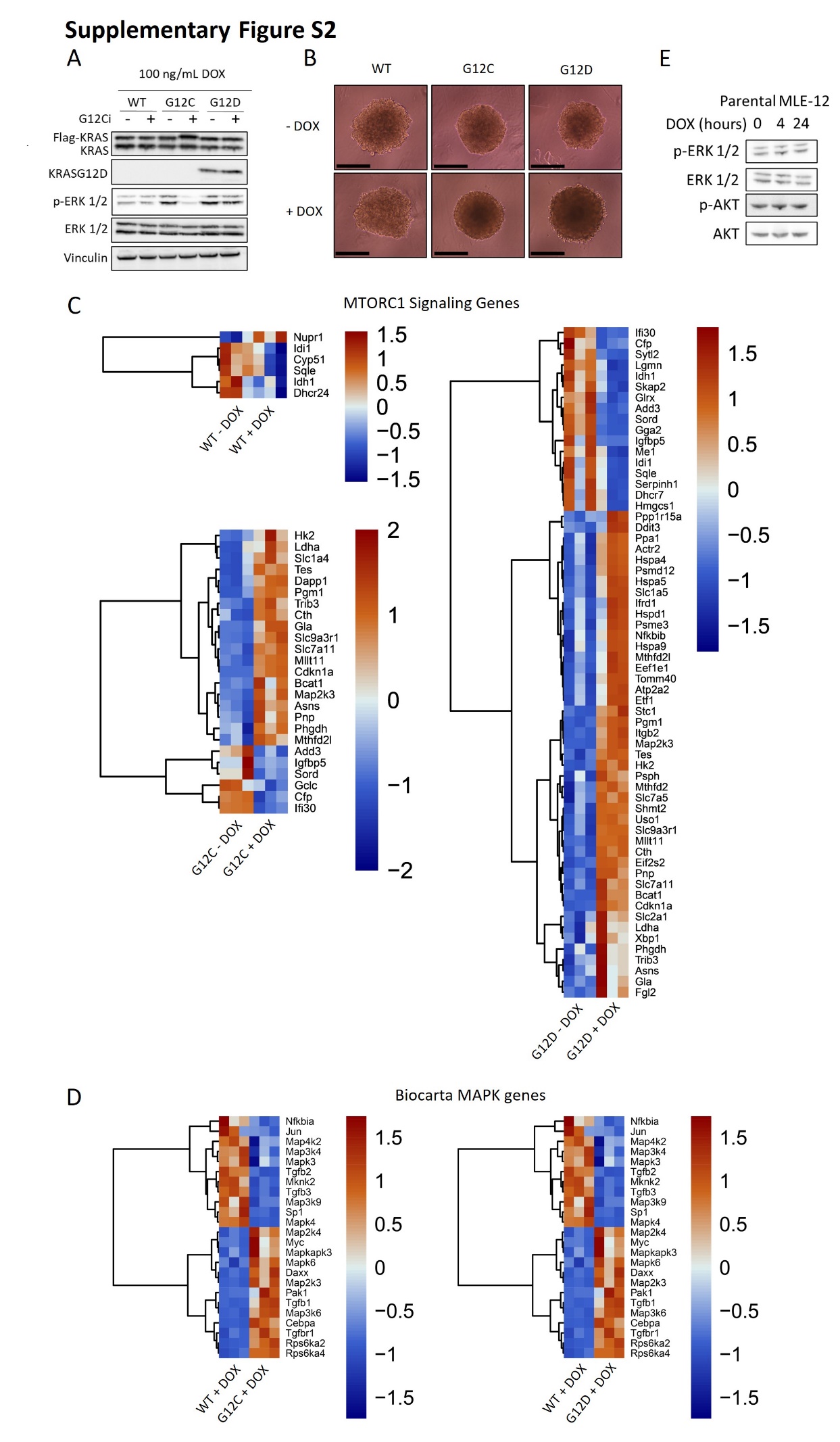

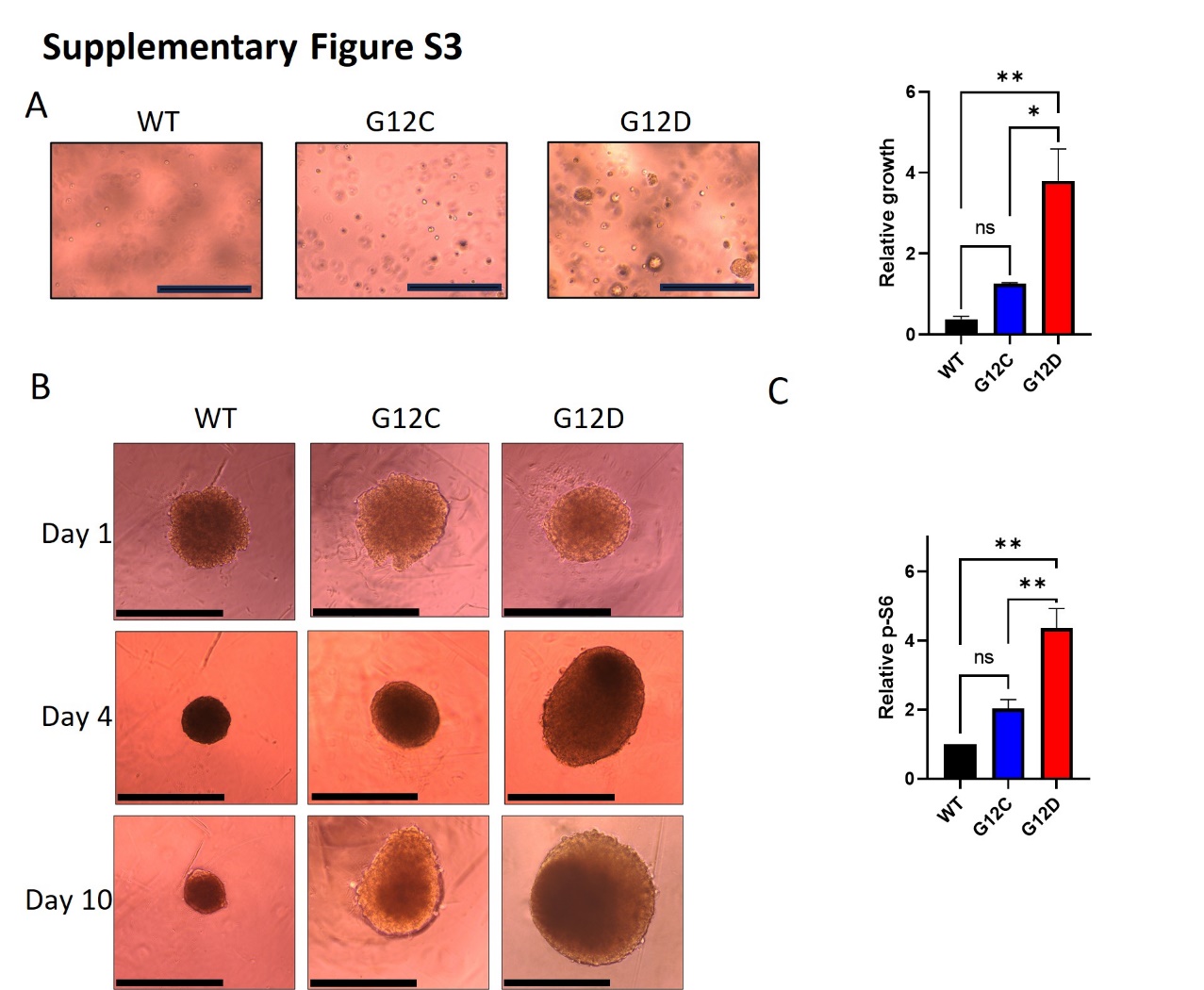

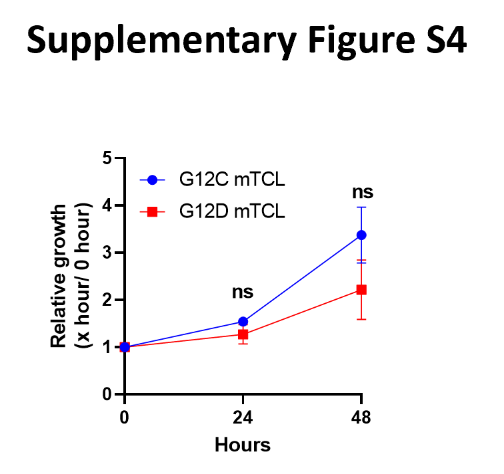

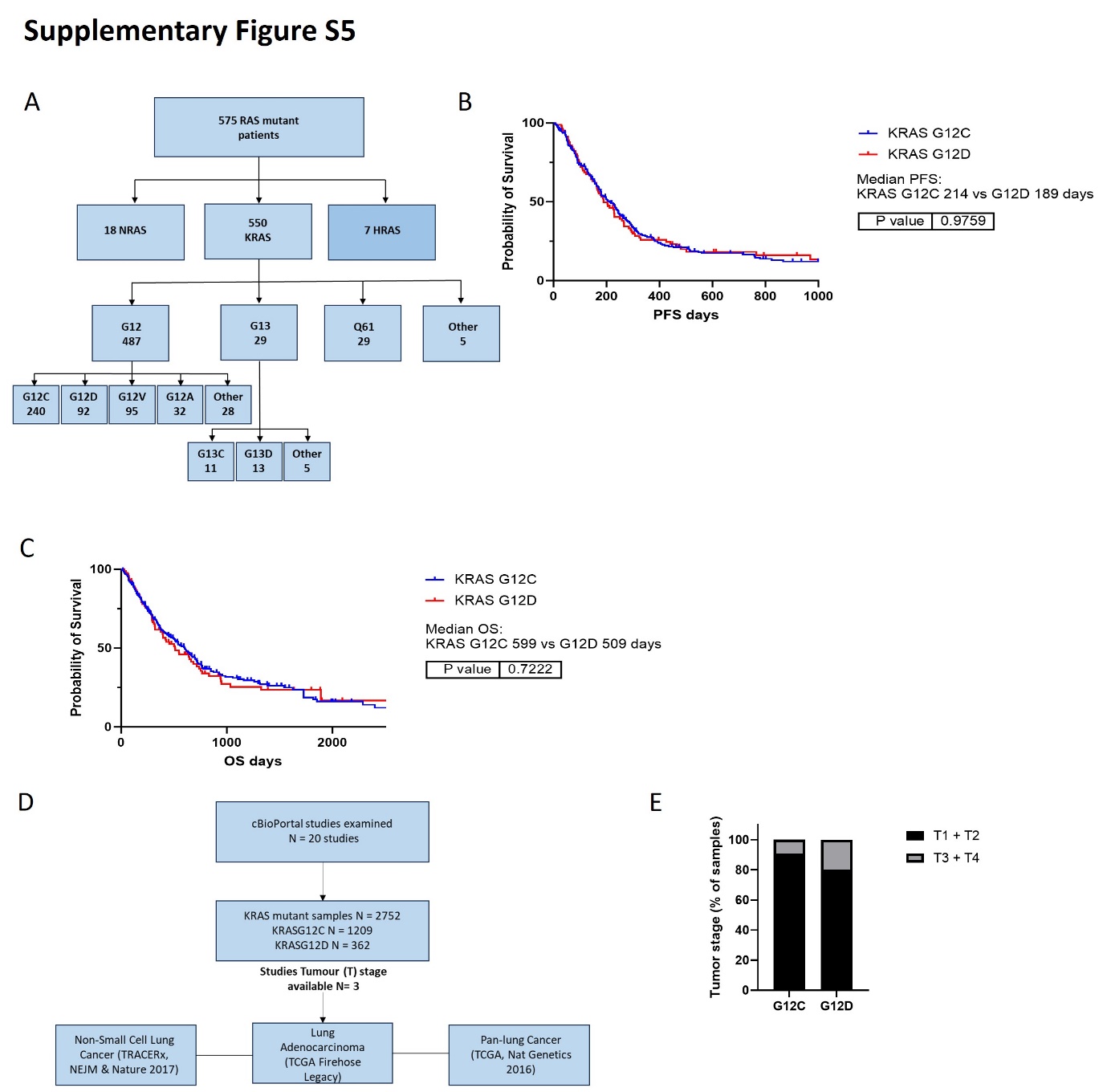

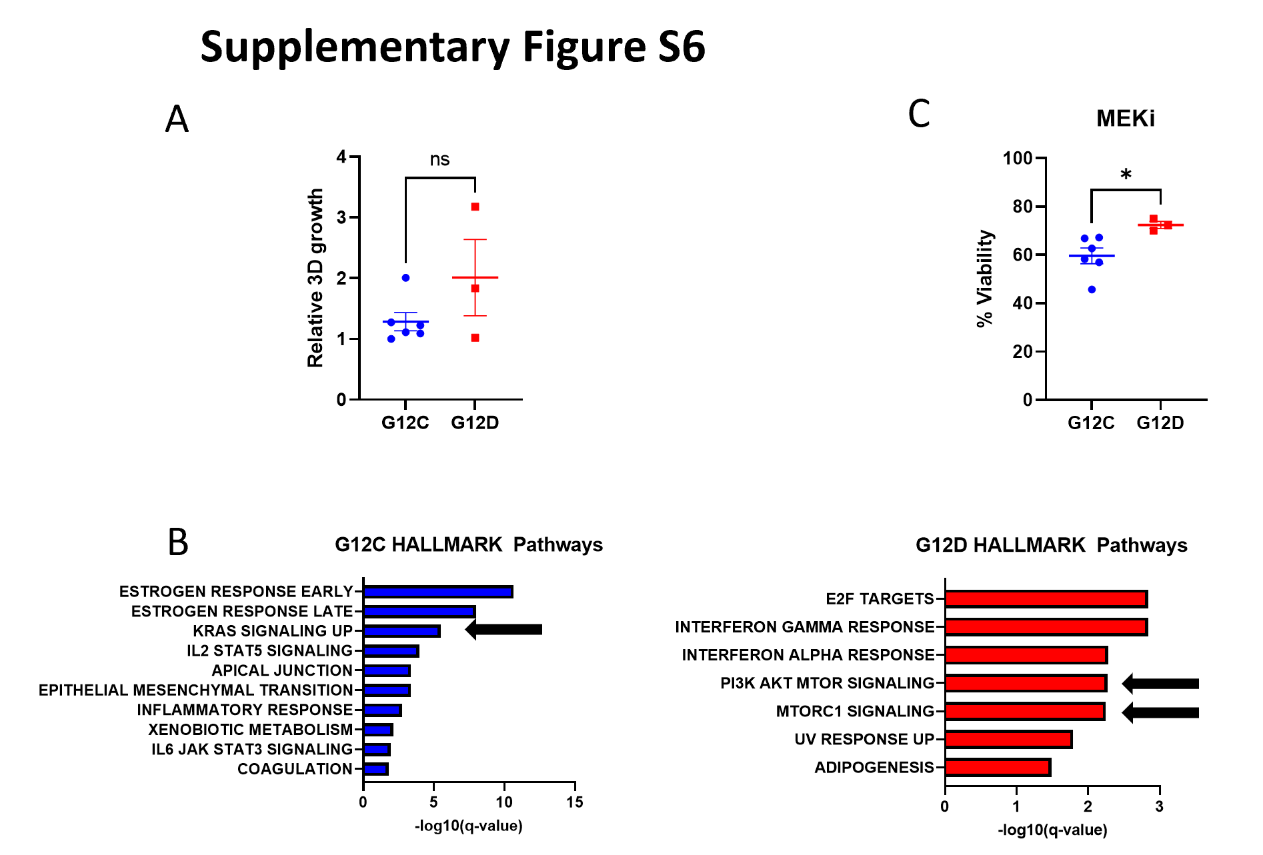

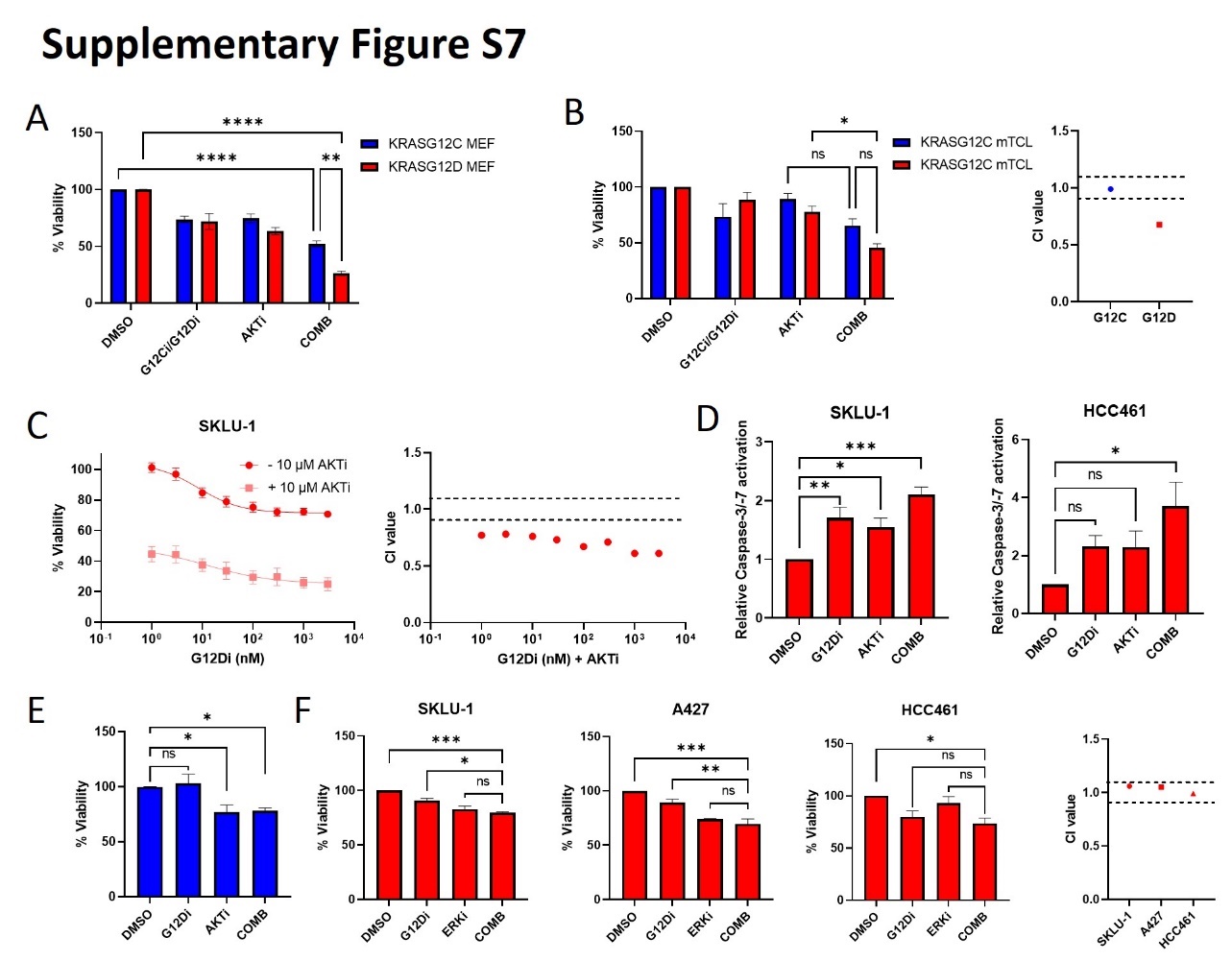

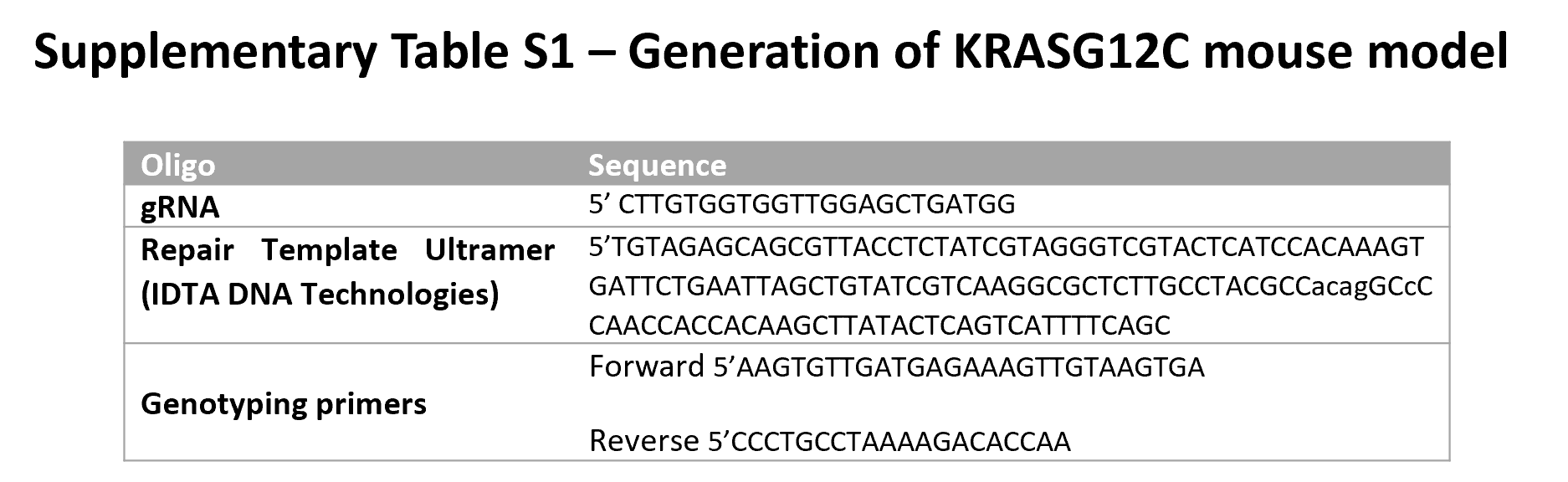

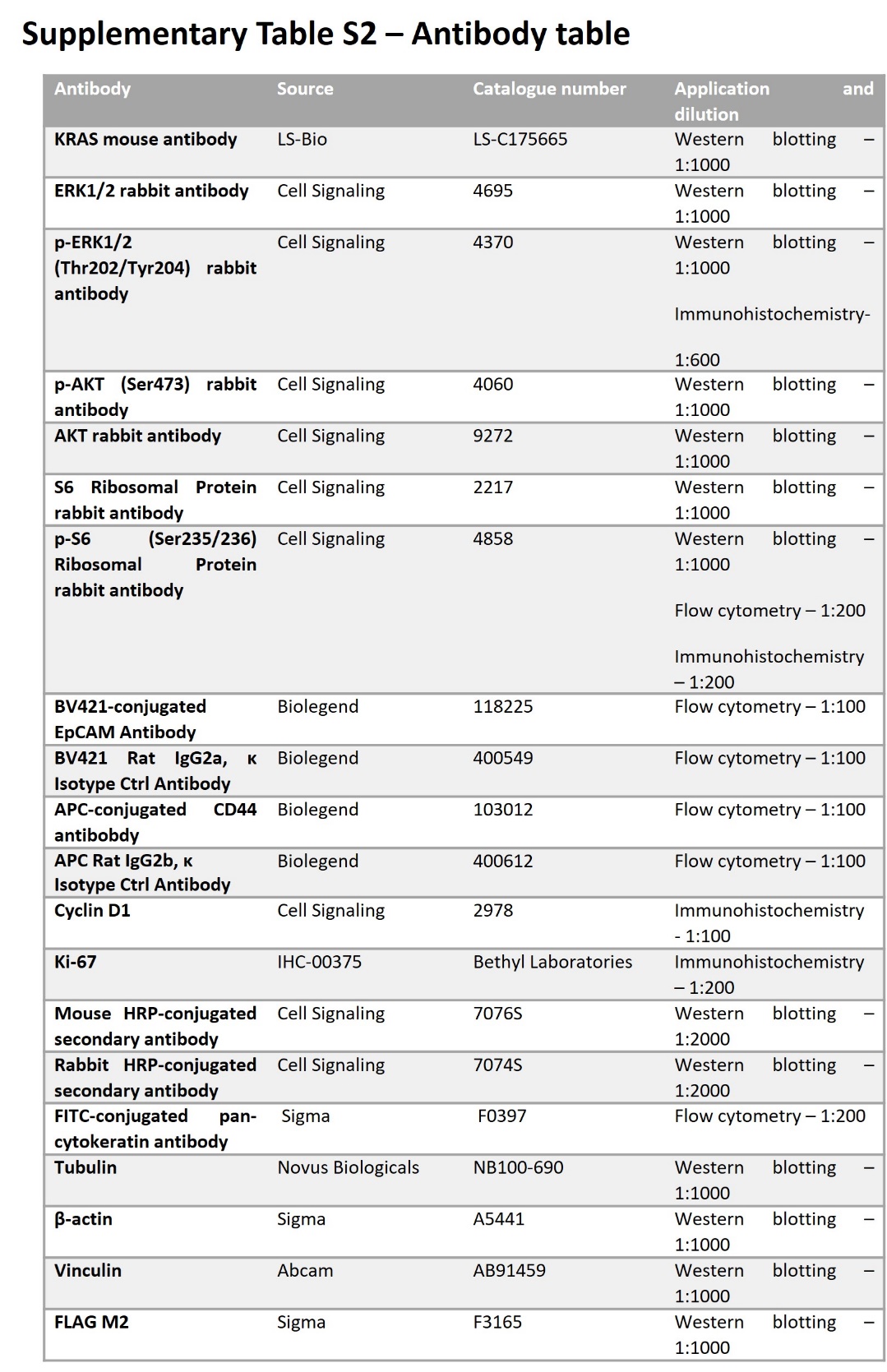

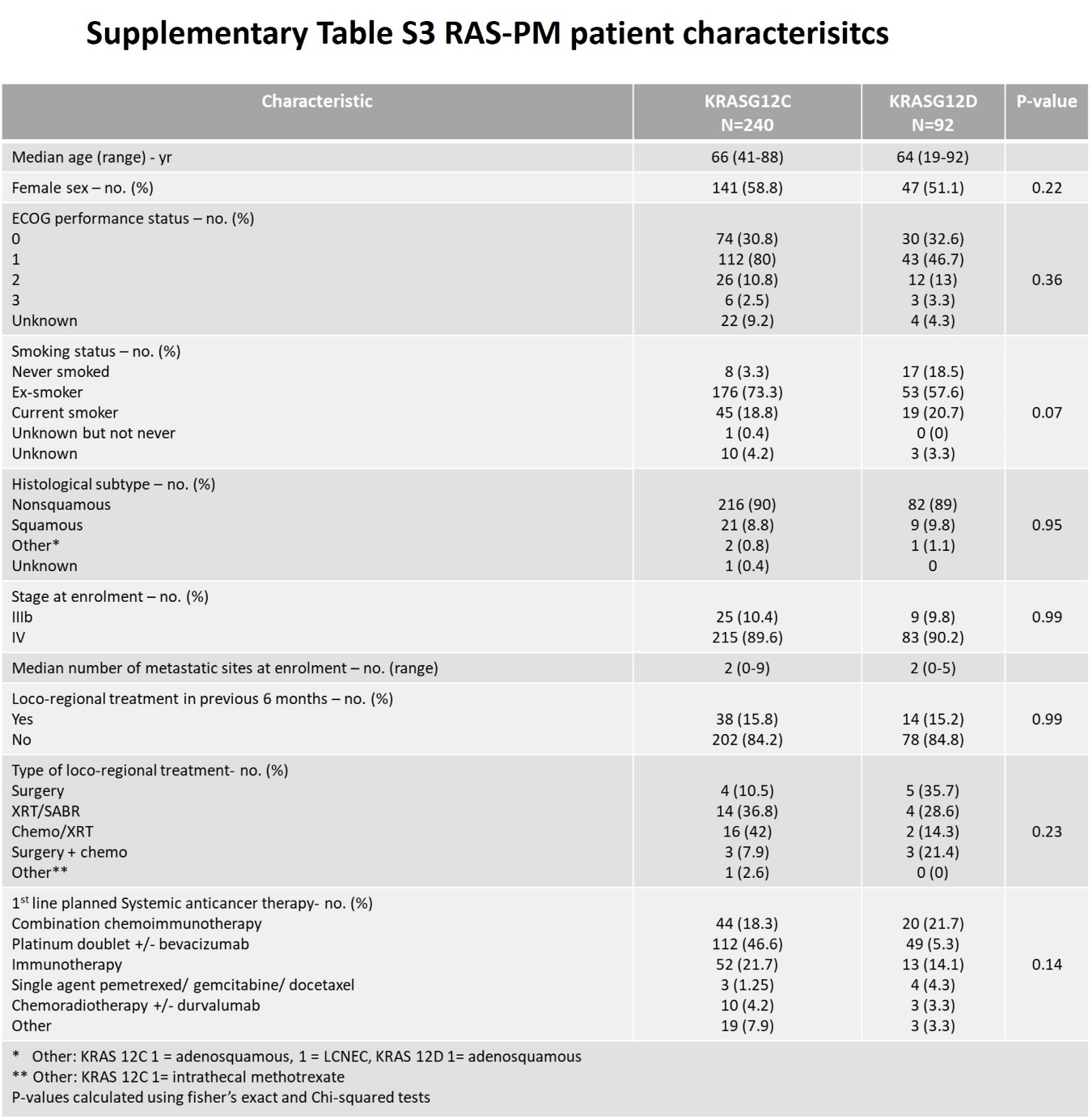
